# A molecularly defined and spatially resolved cell atlas of the whole mouse brain

**DOI:** 10.1101/2023.03.06.531348

**Authors:** Meng Zhang, Xingjie Pan, Won Jung, Aaron Halpern, Stephen W. Eichhorn, Zhiyun Lei, Limor Cohen, Kimberly A. Smith, Bosiljka Tasic, Zizhen Yao, Hongkui Zeng, Xiaowei Zhuang

## Abstract

In mammalian brains, tens of millions to billions of cells form complex interaction networks to enable a wide range of functions. The enormous diversity and intricate organization of cells in the brain have so far hindered our understanding of the molecular and cellular basis of its functions. Recent advances in spatially resolved single-cell transcriptomics have allowed systematic mapping of the spatial organization of molecularly defined cell types in complex tissues^1–3^. However, these approaches have only been applied to a few brain regions^1–11^ and a comprehensive cell atlas of the whole brain is still missing. Here, we imaged a panel of >1,100 genes in ∼8 million cells across the entire adult mouse brain using multiplexed error-robust fluorescence in situ hybridization (MERFISH)^12^ and performed spatially resolved, single-cell expression profiling at the whole-transcriptome scale by integrating MERFISH and single-cell RNA-sequencing (scRNA-seq) data. Using this approach, we generated a comprehensive cell atlas of >5,000 transcriptionally distinct cell clusters, belonging to ∼300 major cell types, in the whole mouse brain with high molecular and spatial resolution. Registration of the MERFISH images to the common coordinate framework (CCF) of the mouse brain further allowed systematic quantifications of the cell composition and organization in individual brain regions defined in the CCF. We further identified spatial modules characterized by distinct cell-type compositions and spatial gradients featuring gradual changes in the gene-expression profiles of cells. Finally, this high-resolution spatial map of cells, with a transcriptome-wide expression profile associated with each cell, allowed us to infer cell-type-specific interactions between several hundred pairs of molecularly defined cell types and predict potential molecular (ligand-receptor) basis and functional implications of these cell-cell interactions. These results provide rich insights into the molecular and cellular architecture of the brain and a valuable resource for future functional investigations of neural circuits and their dysfunction in diseases.

## Introduction

Mammalian brain functions are orchestrated by coordinated actions and interactions of many specialized cell types, including several major classes of cells, such as neurons, glial cells, vascular cells, and immune cells, and numerous distinct cell types within each class. The distinct behaviors and functions of different types of cells are, in a large part, determined by their different molecular properties. Hence, single-cell RNA sequencing (scRNA-seq) provides a systematic approach to classify cell types through gene-expression profiling of individual cells^13–16^. Single-cell epigenomic profiling further enables systematic characterizations of gene-regulatory signatures of different cell types^16–19^. Indeed, numerous molecularly distinct cell types have been identified in the mammalian brain using scRNA-seq and single-cell epigenomic sequencing (for example, Refs.^20–41^). For example, several hundred transcriptionally distinct cell populations have been identified across the entire mouse brain through scRNA-seq of ∼500,000 to 700,000 cells^25, 26^. Despite being a heroic effort at the time, the limited sampling sizes in these studies likely led to an underestimation of the cellular diversity of the brain. Indeed, a recent effort by the BRAIN Initiative Cell Census Network (BICCN) identified ∼100 molecularly distinct cell populations in the mouse primary motor cortex^16^, a small brain region that occupies only a few percent of the total brain volume. It is thus possible that the whole mouse brain contains thousands of molecularly distinct cell populations.

Moreover, understanding the molecular and cellular mechanisms underlying brain functions requires not only a comprehensive classification of cells and their molecular signatures, but also a detailed characterization of how these cells are spatially organized and how they interact with each other. The brain is made of several major regions, including the olfactory areas, isocortex, hippocampal formation, cortical subplate, striatum, pallidum, thalamus, hypothalamus, midbrain, hindbrain, and cerebellum. Each major region further comprises sub-structures that have distinct cell compositions and perform distinct functions. For example, the cerebral cortex forms layered structures, and information is processed by different cortical layers that contain different cell types^42–44^, whereas in subcortical regions, such as thalamus and hypothalamus, neurons often organize into nuclei, which could be structural and functional units for behavior control^45–48^. At a finer scale, spatial location is also a major determinant of cell-cell interactions and communications. While synaptic communications can occur between neurons whose cell bodies are far apart, interactions between neurons and non-neuronal cells, as well as among non-neuronal cells, often occur through direct soma contact or paracrine signaling and hence require spatial proximity between cells. In addition, interactions involving local interneurons also tend to occur between spatially proximal neurons. Therefore, a high-resolution, spatially resolved cell atlas of the brain would provide a valuable resource and reference for understanding the molecular and cellular basis of brain function. Recent advances in spatially resolved transcriptomics have enabled gene-expression profiling and cell-type identification while maintaining the spatial information of cells in intact tissues^1, 2^. These approaches have been used to generate spatial atlases of molecularly defined cell types for a few regions in the mouse and human brain (for example, Refs.^1–11^). However, a high-resolution cell atlas of the whole brain is still missing.

Here, we used a single-cell transcriptome imaging method, multiplexed error-robust fluorescence in situ hybridization (MERFISH)^12^, to generate a molecularly defined and spatially resolved cell atlas of the entire adult mouse brain. By imaging ∼8 million cells across the adult mouse brain and integrating the whole-brain MERFISH and scRNA-seq data, we determined the spatial organization of >5,000 transcriptionally distinct cell clusters, belonging to ∼300 cell subclasses, across the whole mouse brain. This integration also allowed us to impute a transcriptome-wide expression profile for each cell imaged by MERFISH. We further registered the spatial cell atlas generated by MERFISH to the Allen Mouse Brain Common Coordinate Framework (CCF)^49^, providing a reference cell atlas that can be broadly used by the community. This CCF registration further allowed us to qualify the cell-type composition and spatial organization of individual brain regions. Finally, using spatial proximity and ligand-receptor co-expression analyses, we predicted interactions or communications between several hundred pairs of cell types (at the subclass level), and determined ligand-receptor pairs, as well as other genes, that were upregulated in spatially proximal cell pairs, providing insights into potential molecular mechanisms and functional implications of these predicted cell-cell interactions.

### MERFISH imaging of the whole mouse brain

To perform spatially resolved single-cell transcriptomic profiling of the whole mouse brain, we selected a panel of >1,100 genes for MERFISH imaging (**Supplementary Table 1**) based on a whole-brain scRNA-seq dataset (∼4 million cells) described in a companion manuscript in this BICCN package (Yao et al.). Clustering analysis of the scRNA-seq data resulted in 5,200 transcriptomically distinct cell clusters, which were grouped into 306 subclasses (Yao et al.). Our MERFISH gene panel was selected from marker genes differentially expressed between these subclasses and clusters, comprising 23 neurotransmitter-related genes, 21 neuropeptide genes, 187 transcription factor genes, 123 subclass markers (partially overlapping with some of the above-mentioned genes), as well as other genes differentially expressed between pairs of cell clusters (see **Methods** for details) (**Figure 1a**).

**Figure 1.**
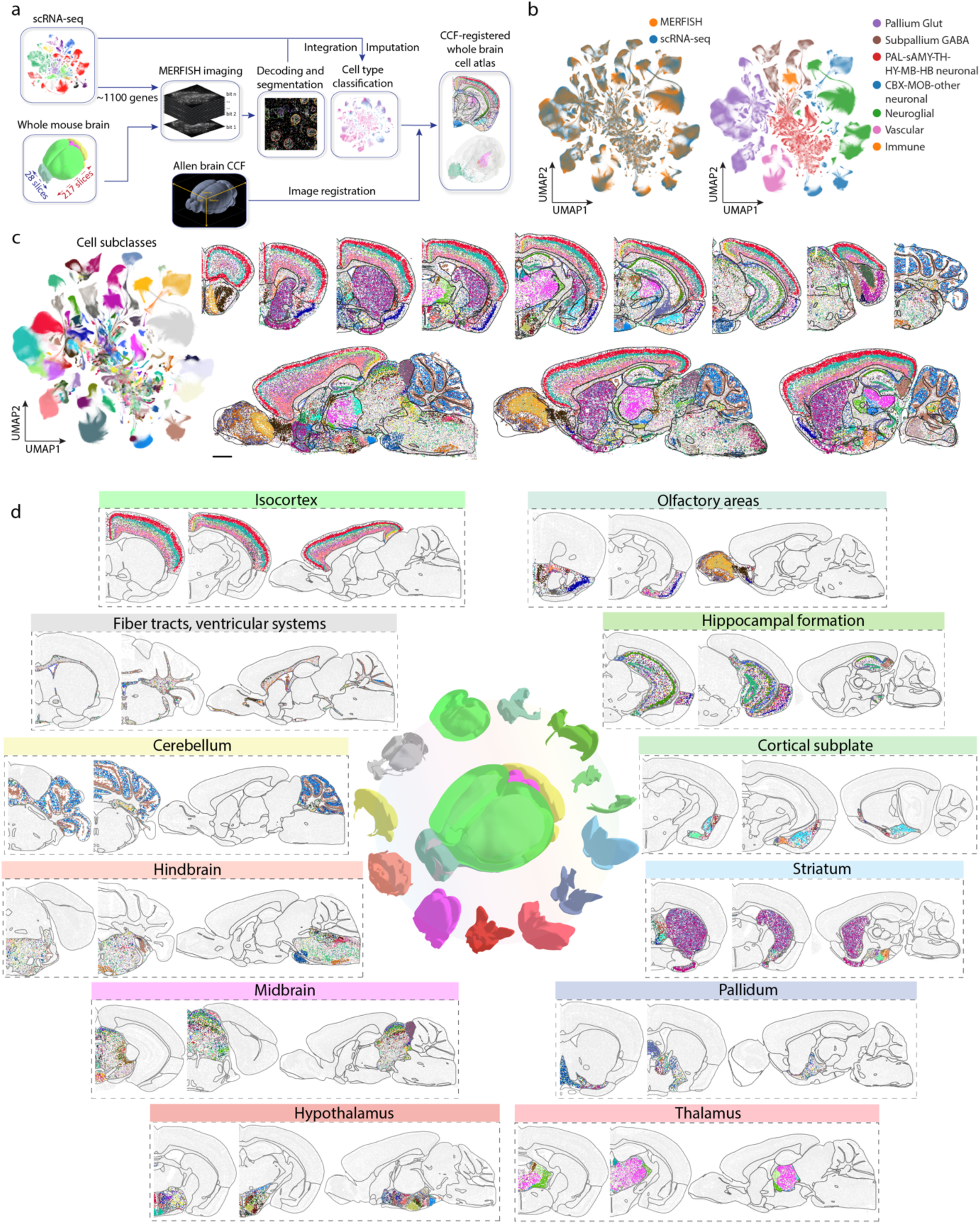
A molecularly defined and spatially resolved cell atlas of the whole mouse brain. **a**, Workflow to construct a spatially resolved whole mouse brain cell atlas. A panel of genes were chosen based on the clustering results from scRNA-seq data and were used for MERFISH imaging. Whole adult mouse brains were sliced to generate serial coronal or sagittal sections for MERFISH imaging. The MERFISH images were decoded and segmented, and the resulting single-cell gene expression profiles were integrated with the scRNA-seq data to classify cells in the MERFISH datasets and impute transcriptome-wide expression profiles for each imaged cell. The MERFISH images were then registered to the Allen CCF version 3 to create a spatial atlas of molecularly defined cell types across the whole mouse brain in the CCF space. **b**, Uniform manifold approximation and projection (UMAP) of the integrated scRNA-seq and MERFISH data with cells coloured by measurement modalities (left) or by seven major divisions of cells (right). The number of cells in the MERFISH or scRNA-seq dataset in each of the 306 subclasses was downsampled to the corresponding number in the other dataset for visualization purpose, such that one color does not dominate the other in the left panel. The integration UMAP with all MERFISH and scRNA-seq cells displayed is shown in **Extended Data Figure 1d**. **c**, Left: UMAP of the integrated MERFISH and scRNA-seq data. Right: Spatial maps of the cell types in example coronal and sagittal sections. Cells are coloured by their subclass identities in the UMAP and spatial maps. The black lines in the spatial maps mark the major brain region boundaries defined in the CCF. Scale bar: 1 mm. In this and subsequent figures, all cells are shown in the experimental coordinates and the boundaries of brain regions defined in the CCF were transformed to the experimental coordinates based on our CCF registration results (See **Methods**). **d**, Spatial maps of the cell types in example coronal and sagittal sections in the 11 major brain regions as well as in fiber tracts and ventricular systems. Cells are coloured by their subclass identities as in **c**.

We imaged these genes in a series of 10-μm-thick coronal and sagittal sections spanning whole hemispheres of the adult mouse brain, including serial coronal sections at 100-μm intervals (Animal 2, female, 150 sections) or 200-μm intervals (Animal 1, male, 67 sections), and serial sagittal sections at 200-μm intervals (Animals 3 and 4, male, 28 sections total with only 3 sections from Animal 4 to compensate for the broken sections from Animal 3) (**Figure 1a**). Individual RNA molecules were identified and assigned to individual cells segmented based on the DAPI and total RNA signals, providing the expression profiles of individual cells (see **Methods**). The MERFISH data exhibited excellent reproducibility between replicate animals (**Extended Data Figure 1a**). The mean copy number per cell for individual genes obtained from MERFISH measurements correlated well with the mean expression levels determined from whole-brain bulk RNA-seq (**Extended Data Figure 1b**) and scRNA-seq data (**Extended Data Figure 1c**).

In total we imaged ∼8 million cells across the adult mouse brain, including all 11 major brain regions: olfactory areas (OLF), isocortex (CTX), hippocampal formation (HPF), cortical subplate (CTXsp), striatum (STR), pallidum (PAL), thalamus (TH), hypothalamus (HY), midbrain (MB), hindbrain (HB), and cerebellum (CB).

### Cell classification and registration to the common coordinate framework

In order to classify the imaged cells, we integrated the MERFISH data with the scRNA-seq data using a canonical correlation analysis (CCA)-based integration method^35, 50^ and classified each MERFISH cell based on the most frequently appearing cell-type identity among the 100 nearest-neighbour anchor cells in the scRNA-seq dataset in the integrated gene-expression space (see **Methods**, **Figure 1a**). The MERFISH and scRNA-seq data integrated well with each other (**Figure 1b**, **Extended Data Figure 1d**), and the cell-type labels were transferred from the scRNA-seq cells to the MERFISH cells with high confidence scores (see **Methods**, **Extended Data Figure 1e**). We set a threshold on the confidence scores for cell-type label transfer (0.8 for subclass label transfer: >80% of the 100 nearest-neighbour anchor scRNA-seq cells must have the same subclass label for this label to be transferred to a MERFISH cell; 0.5 for cluster label transfer: >50% of the 100 nearest-neighbour anchor cells must bear the same cluster label for label transfer to occur). 82% and 75% MERFISH cells passed the subclass and cluster confidence score thresholds, respectively, and were used for subsequent analysis. To further test the robustness of label transfer, we performed label transfer with an alternative approach by calculating the cosine distances of the gene expression profiles between each MERFISH cell and the scRNA-seq clusters and assigning each MERFISH cell with the label of the closest scRNA-seq cluster. Results from these two methods showed excellent agreement (**Extended Data Figure 1f**). Overall, all 306 subclasses and 99% (5,139) of the 5,200 clusters identified by scRNA-seq were observed in the MERFISH data.

Integration of the MERFISH and scRNA-seq data also allowed us to impute the transcriptome-wide expression profile for the MERFISH-imaged cells. Specifically, for each MERFISH cell, we computed the weighted average expression profile of the 30 nearest-neighbour anchor cells in the scRNA-seq dataset and assigned this average expression file to the MERFISH cell. To validate the imputation results, for the genes in the MERFISH gene panel, we compared the imputed gene expression levels with the values directly measured by MERFISH and the previously measured spatial expression patterns in the Allen Brain Atlas in situ hybridization data^51^; for the genes that were not included the MERFISH gene panel, we compared the spatial patterns determined from the imputation results with the Allen Brain Atlas in situ hybridization data^51^. In both cases, we obtained excellent agreement (**Extended Data Figure 2**).

To enable systematic quantifications of the cell composition and organization in different brain regions, we registered the cell atlas generated by MERFISH to the Allen mouse brain CCF version 3 (http://atlas.brain-map.org/)49 (**Figure 1a**) using a two-step procedure, in which we first aligned the DAPI images in the MERFISH dataset to the Nissl template images in the Allen Reference Atlas and then refined the alignment with cell-type-based landmarks (see **Methods**, **Extended Data Figure 3a**). This CCF registration allowed us to place each individual MERFISH-imaged cell, with the cell-type-identity label, into the 3D common reference space (**Figure 1c, d; Extended Data Figure 3b**).

The spatial location information of the cell subclasses measured by MERFISH were also used for the annotation of the cell subclasses identified by scRNA-seq, as described in the companion manuscript (Yao et al.). Briefly, except for some of the previously well-annotated subclasses, each neuronal subclass name has three parts: the brain region in which the subclass primarily resides (e.g., L2/3, MEA-BST, LSX, etc.), one or more major marker genes (e.g., *Pmch*, *Tfap2b*, *Prdm12*, etc.), and the major neurotransmitter (e.g., Glut, Gaba, Dopa, etc.) expressed in the subclass. For example, “LSX Prdm12 do Gaba” stands for the GABAergic neuronal subclass marked by *Prdm12* residing in the dorsal (do) part of the lateral septal complex (LSX) in the striatum. Non-neuronal cell subclasses were annotated primarily based on marker genes and named based on prior knowledge (for example, Microglia, Astrocyte, etc.) with spatial information being specified only in some cases (for example, Astro-OLF for an astrocyte subclass residing in olfactory areas). For both neurons and non-neuronal cells, the cell clusters were named by the subclass names followed by numerical indices.

### Cellular diversity and spatial organization of neurons

Registration of the MERFISH images to the Allen CCF allowed us to quantify the composition and organization of cell types in individual brain regions (**Figure 1d**). Overall, the whole mouse brain consisted of 43% neurons and 57% non-neuronal cells. This ratio varied substantially from region to region, with hindbrain and cerebellum showing the lowest and highest neuronal-to- non-neuronal cell ratio, respectively (**Figure 2a**).

**Figure 2.**
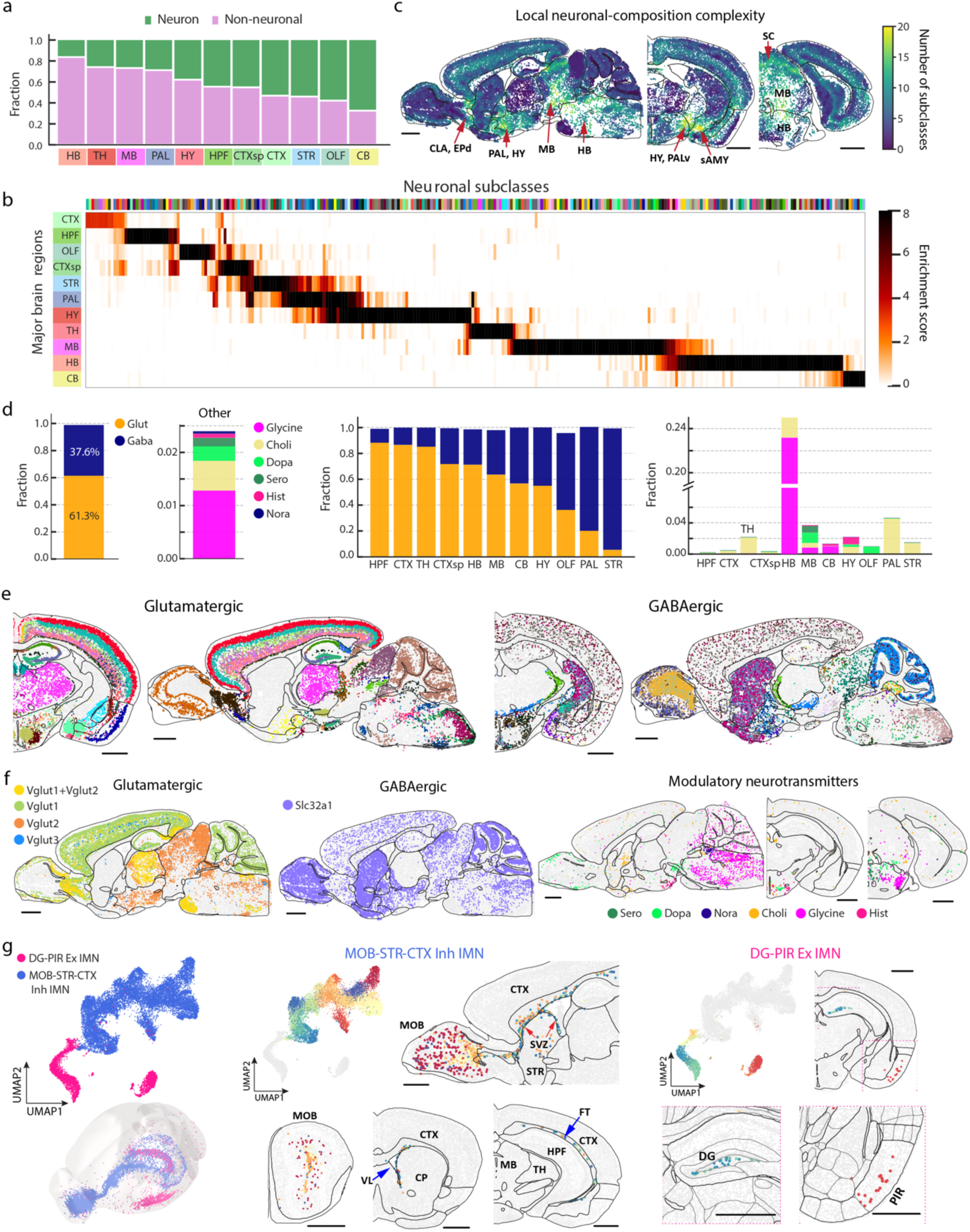
Cell compositions and spatial distributions of neurons across the whole brain. **a**, Fractions of neurons and non-neuronal cells in the 11 major brain regions. **b**, Heatmap showing the enrichment score of each neuronal subclass in the 11 major brain regions. The coloured bars at the top and on the left indicate the neuronal subclasses and brain regions, respectively. The enrichment score of a subclass in each individual brain regions is defined as the fold change of the average cell density of a subclass within a brain region compared to the average density across the whole brain. **c**, Spatial heatmap of local neuronal-composition complexity. The local complexity of neuronal cell-type composition in the neighbourhood of any given cell is defined as the number of different neuronal cell types (at the subclass level) present in the 50 nearest-neighbour neurons surrounding that cell. **d**, Left two panels: Bar plots showing the fractions of neurons using different neuronal transmitters across the whole brain. Right two panels: The fractions of neurons using different neuronal transmitters in individual brain regions. Glut: glutamatergic neurons; Gaba: GABAergic neurons; Glycine: glycinergic neurons; Choli: cholinergic neurons; Dopa: dopaminergic neurons; Sero: serotonergic neurons; Hist: histaminergic neurons; Nora: Noradrenergic neurons. **e**, Spatial maps of the glutamatergic (left) and GABAergic (right) neuronal subclasses in one coronal section and one sagittal section as examples, with cells coloured by their subclass identities. **f**, Spatial maps of the glutamatergic neurons expressing *Vglut1*, *Vglut2*, *Vglut1*+*Vglut2*, and *Vglut3* (left), the GABAergic neurons (middle), and the neurons expressing various modulatory neurotransmitters (right). *Vglut3* expressing neurons often co-express *Vglut1* and/or *Vglut2*. Neurons expressing different neurotransmitters are identified by the expression of transporters or synthesis enzymes of these neurotransmitters. **g**, Left: UMAP (top) and spatial distribution (bottom) of the immature neurons (IMNs) with neurons coloured by subclass identities (pink: excitatory IMNs; blue: inhibitory IMNs). Spatial distributions are shown in the 3D CCF space. Middle: UMAP (top left) and spatial distribution (other panels) of the inhibitory IMNs shown in sagittal and coronal sections coloured by cluster identities of the IMNs. Excitatory IMNs are shown in grey in the UMAP. Right: UMAP (top left) and spatial distribution of the excitatory IMNs shown in a coronal section coloured by cluster identities (top right). Inhibitory IMNs are shown in grey in the UMAP. Zoomed-in views of the two distinct locations of excitatory IMNs in the dentate gyrus (DG) area and the piriform area (PIR) are shown in the bottom panels. Scale bars in **c, e-g**: 1 mm.

Neurons exhibited an exceptionally high level of diversity. Among the 306 subclasses and 5,200 clusters identified, 283 subclasses and >5,000 clusters were neurons. The spatial distributions of the neuronal cell types showed strong regional specificity in the brain. To obtain a systematic picture of where different neuronal cell types were located, we calculated an enrichment score of each neuronal subclass across the 11 major brain regions by comparing the average cell density of a subclass within a region to that across the whole brain. Most neuronal subclasses were only enriched in one single major region, while some subclasses spanned multiple, usually physically connected, regions, such as striatum and pallidum, pallidum and hypothalamus, or midbrain and hindbrain (**Figure 2b**).

Many of the subclass boundaries aligned well with the region boundaries shown in the CCF. For example, the IT (intratelencephalic) subclasses showed a clean separation at the boundaries between isocortex and olfactory areas or hippocampal formation (**Extended Data Figure 4a**). In thalamus, AV Col27a1 Glut and AD Serpinb7 Glut perfectly fit in anteroventral (AV) and anterodorsal (AD) nucleus, respectively (**Extended Data Figure 4b**). In the meantime, we also observed some subclasses spanning multiple brain regions. For example, inhibitory neuronal subclasses marked by *Lamp5*, *Sncg*, *Vip*, *Sst*, or *Pvalb* were distributed across isocortex, hippocampal formation, olfactory areas, and cortical subplate (**Extended Data Figure 4c**), consistent with previous knowledge^30, 52^.

For each of the 11 major regions, we quantified their cell-type composition at the subclass and cluster level (**Supplementary Table 2**). The number of cell types that were contained in each brain region differ from region to region. In particular, midbrain, hindbrain, and hypothalamus regions contained substantially greater number of neuronal cell types compared to the other brain regions (**Figure 2b**). We further quantified the local complexity of neuronal cell-type composition, defined as the number of distinct neuronal cell types (subclasses) present in the neighbourhood of each cell (neighbourhood: 50 nearest-neighbour neuronal cells). Notably, the local complexity of neuronal cell-type composition was also substantially higher in midbrain, hindbrain, and hypothalamus, as compared to other major brain regions (**Figure 2c**), indicating that these regions were not simply composed of more subregions with simple cell compositions, but each local neighbourhood within these regions also tended to have higher cellular diversity. In addition, a few subregions in some other main regions such as the claustrum (CLA) and endopiriform nucleus (EP) in the cortical subplate and the hypothalamus-neighbouring regions such as the striatum-like amygdalar nuclei (sAMY) and ventral pallidum (PALv) also exhibited a high local complexity of neuronal cell-type composition (**Figure 2c**).

### Spatially dependent neurotransmitter and neuropeptide usage of neurons

Next, we examined the neurotransmitter usage of neurons in different brain regions. Based on the expression of neurotransmitter transporters and genes involved in neurotransmitter biosynthesis, we classified matured neurons into eight partially overlapping groups: glutamatergic (expressing *Slc17a7*, *Slc17a6* and/or *Slc17a8*), GABAergic (expressing *Slc32a1*), serotonergic (expressing *Slc6a4*), dopaminergic (expressing *Slc6a3*), cholinergic (expressing *Slc18a3*), glycinergic (expressing *Slc6a5*), noradrenergic (expressing *Slc6a2*), and histaminergic (expressing *Hdc*) neurons.

Among these, glutamatergic and GABAergic neurons accounted for ∼61% and ∼38% of the total neuronal populations, respectively, whereas serotonergic, dopaminergic, cholinergic, glycinergic, noradrenergic, and histaminergic neurons (often co-expressing glutamate or GABA transporters) accounted for only ∼2-3% of the total neuronal population (**Figure 2d, left**). Both glutamatergic and GABAergic neurons were widely distributed across the whole brain and were classified into diverse cell types with distinct spatial distributions across different brain regions (**Figure 2e, f**). The glutamatergic-to-GABAergic neuron ratio (Excitatory:Inhibitory balance) varied drastically from brain region to brain region (**Figure 2d**, **middle**). Among the 11 major brain regions, hippocampal formation, isocortex, and thalamus had the highest glutamatergic-to-GABAergic neuron ratio, ∼6:1 – 8:1, whereas this ratio was the lowest (∼1:14) in striatum, which was dominated by the GABAergic medium spiny neurons (MSNs). Although thalamus was mostly made of glutamatergic neurons, the reticular nucleus (RT) of thalamus was dominated by GABAergic neurons (**Extended Data Figure 4d**). GABAergic neurons also dominated in pallidum. In midbrain and hindbrain, glutamatergic and GABAergic neurons were widely distributed in a partially intermingled manner (**Extended Data Figure 4e**). In cerebellum, glutamatergic and GABAergic neurons were separately enriched in the granular and molecular layers, respectively, as expected (**Extended Data Figure 4f**). A small fraction of neurons (∼1%) exhibited co-expression of both glutamate and GABA neurotransmitter transporters (*Slc17a6/7/8* and *Slc32a1*, respectively) and these neurons were primarily found in non-telencephalic areas of the brain such as the globus pallidus internal segment (GPi), hypothalamic nuclei such as the anterior hypothalamic nucleus (AHN) and supramammillary nucleus (SUM), and some subregions in midbrain and hindbrain, as well as in the outer layer of the main olfactory bulb (MOB) (**Extended Data Figure 4g**), both corroborating and expanding previously knowledge that neurons co-releasing glutamate and GABA are present in GPi and hypothalamus^4, 53–56^.

Among the glutamatergic neurons, *Vglut1* (*Slc17a7*), *Vglut2* (*Slc17a6*), and *Vglut3* (*Slc17a8*) were differentially distributed in different brain regions (**Figure 2f, left**)^57^. *Vglut1* dominated in olfactory areas, isocortex, hippocampal formation, cortical subplate, as well as in the cerebellar cortex, whereas *Vglut2* dominated in hypothalamus, midbrain, and hindbrain. In some regions, *Vglut1* and *Vglut2* were co-expressed in neurons, such as the retrosplenial areas (RSP), pontine gray (PG), anterior olfactory nucleus (AON), and thalamus (**Figure 2f, left**, **Extended Data Figure 4h**). The less used *Vglut3* were scattered across multiple brain regions, enriched in regions such as layer 5 of isocortex and bed nuclei of the stria terminalis (BST), and were often co-expressed with *Vglut1* and/or *Vglut2* (**Figure 2f, left**).

We also located the neurons that used other, modulatory neurotransmitters (**Figure 2f, right**). Dopaminergic neurons were observed in olfactory areas (located in the glomerular layer), hypothalamus (enriched in the arcuate hypothalamic nucleus (ARH)), and midbrain (enriched in the ventral tegmental area (VTA) and neighbouring areas) (**Extended Data Figure 4i**)^58^. Serotonergic neurons were enriched in the raphe nuclei (DR, RPO, RM) in midbrain and hindbrain (**Extended Data Figure 4j**)^59^. Histaminergic neurons were observed in ventral tuberomammillary nucleus (TMv), tuberal nucleus (TU), and other neighbouring areas in the ventral hypothalamus (**Extended Data Figure 4k**)^60^. Glycinergic neurons were widely distributed across hindbrain (**Extended Data Figure 4I**)^61^. Noradrenergic neurons were localized to the locus ceruleus (LC) and neighbouring areas in hindbrain (**Extended Data Figure 4m**)^62, 63^. Cholinergic neurons were found in many different locations of the brain, including striatum, ventral pallidum, and multiple small subregions such as lateral septal complex (LSX), medial habenula (MH), ARH, pedunculopontine (PPN) and parabigeminal (PBG) nucleus in midbrain, and dorsal motor nucleus of the vagus nerve (DMX) and nucleus of the solitary tract (NTS) in hindbrain (**Extended Data Figure 4n**)^64^.

These modulatory transmitter transporter genes were often found to be co-expressed with glutamate or GABA transporters in individual neurons. For example, dopaminergic neurons in olfactory areas co-expressed *Slc32a1*, and in midbrain and hypothalamus, co-expression with *Slc32a1* or *Slc17a6* were both observed. Cholinergic neurons in striatum and pallidum co-expressed *Slc32a1* and those in hindbrain also co-expressed *Slc17a6*. Glycinergic neurons and histaminergic neurons co-expressed *Slc32a1*.

Our MERFISH images also showed spatially heterogeneous distributions of many neuropeptide genes (**Extended Data Figure 5**). To name just a few examples, *Adcyap1* and *Gal* were enriched in multiple nuclei in hypothalamus; *Penk* was widely expressed in striatum, midbrain and cerebellum, and particularly enriched in striatum; *Prok2* was expressed in olfactory tubercle (OT) and multiple nuclei in hypothalamus; *Tac2* was enriched in BST and multiple nuclei in hypothalamus, striatum, and thalamus. *Trh* was expressed in RT in the thalamus, hypothalamus, cortical amygdalar area posterior part (COAp) in olfactory areas, and inferior olivary complex (IO) in medulla.

In addition to matured neurons, we also observed two subclasses of immature neurons (IMNs), one inhibitory and one excitatory (**Figure 2g, left**). The inhibitory IMNs, composed of 31 clusters, were distributed along the subventricular zone (SVZ), extending to the olfactory bulb through the anterior commissure (**Figure 2g, middle**), consistent with the previous knowledge of adult neurogenesis in the SVZ and migration of neuroblast to the olfactory bulb along the rostral migratory stream (RMS)^65–67^. The excitatory IMNs, composed of 7 clusters, were found in two distinct locations: cluster 5092 was primarily located in the piriform area (PIR) of olfactory areas, while the other clusters were distributed along the dentate gyrus (DG) in hippocampal formation (**Figure 2g, right**) consistent with the previous knowledge of adult neurogenesis in hippocampal formation^68, 69^.

### Cellular diversity and spatial organization of non-neuronal cells

We also examined the spatial organization of non-neuronal cells, comprising 23 subclasses and 99 clusters (**Figure 3a**), and quantified the non-neuronal cell-type composition and enrichment in the 11 major brain regions, as well as in fiber tracts and ventricular systems where non-neuronal cells dominate (**Figure 3b, c**; **Supplementary Table 2**).

**Figure 3.**
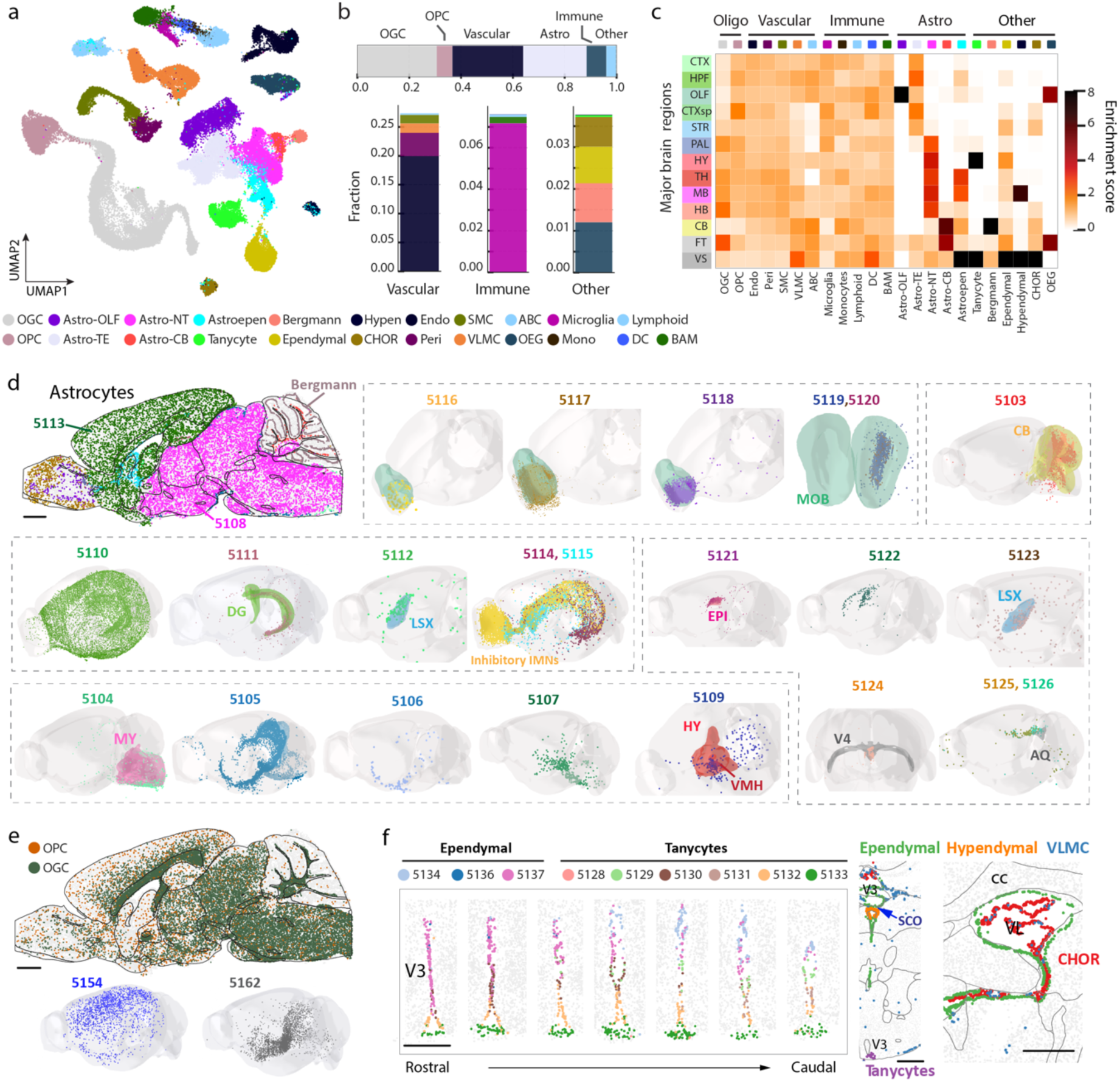
Cell compositions and spatial distributions of non-neuronal cells across the whole brain. **a**, UMAP of non-neuronal cells coloured by their subclass identities as shown in the legend below. **b**, Bar plots showing the fractions of major non-neuronal cell types in the whole brain, including oligodendrocytes, OPC, vascular cells, astrocytes, immune cells, and other cell types (top), and fractions of different vascular cell types, immune cell types, and non-neuronal cell types in the “other” category (bottom). Cell subclasses in the bottom bar plots are coloured as shown in the legend below. OGC: oligodendrocytes; OPC: oligodendrocyte progenitor cell; Endo: endothelial cell; Peri: pericytes; SMC: smooth muscle cell; BAM: border-associated macrophages; DC: dendritic cell; Mono: monocytes; VLMC: vascular lepotomeningeal cell; OEG: olfactory ensheathing glia; CHOR: choroid plexus epithelial cell; ABC: arachnoid barrier cell; Hypen: hypendymal cell. **c**, Heatmap showing the enrichment scores of all non-neuronal subclasses in 11 major brain regions, as well as in fiber tracts (FT) and ventricular systems (VS). The enrichment score is defined as in Figure 2b. **d**, Spatial distributions of the 24 astrocytes clusters and Bergmann cells shown in a sagittal section (top left) and in the 3D CCF space (other panels), with cells coloured by their cluster identities and cluster numerical indices shown. **e**, Spatial distributions of the oligodendrocytes and OPCs shown in a sagittal section with cells coloured by their subclass identities (top). Two specific oligodendrocyte clusters are shown in the 3D CCF space with cells coloured by their cluster identities (bottom). **f**, Left: Spatial maps of three ependymal and six tanycyte clusters in the third ventricle (V3) in seven different coronal sections, 100 μm apart from each other along the rostral-caudal direction. Right: Spatial maps of CHOR, ependymal, hypendymal, and VLMCs in the third ventricle (V3, left) and lateral ventricle (VL, right). Scale bars in **d**, **e**: 1 mm; Scale bars in **f**: 0.5 mm.

Across the whole brain, non-neuronal cells were composed of 31% of oligodendrocytes, 6% of oligodendrocyte progenitor cells (OPC), 27% of vascular cells [endothelial cells, pericytes, vascular leptomeningeal cells (VLMC), smooth muscle cells (SMC), arachnoid barrier cells (ABC)], 24% of astrocytes, 8% of immune cells [microglia, border-associated macrophages (BAM), lymphoid cells, dendritic cells, monocytes], and 4% other cell types [olfactory ensheathing glia (OEG), Bergmann cells, ependymal cells, CHOR cells, tanycytes, hypendymal cells] (**Figure 3b**).

Notably, some of the non-neuronal cell types also exhibited strong regional specificity (**Figure 3c**). Such spatial heterogeneity was particularly pronounced for astrocytes, as well as cells belonging to the ventricular systems (**Figure 3c**). We observed a high diversity of astrocytes, including 24 cell clusters, all of which exhibited regional specificity. Among these, the two biggest clusters, Astro 5113 and Astro 5108, accounting for 47% and 39% of the total astrocyte population, respectively, showed distinct spatial distributions with the former being exclusively located in the telencephalon and the latter in non-telencephalic regions (**Figure 3d**), consistent with previous observations^25^. In addition, Astro clusters 5116-5120 were located in the olfactory bulb; Astro 5103 was located in the cerebellum; Astro 5111 was located in dentate gyrus, Astro 5109 was enriched in the hypothalamus, Astro 5104 was enriched in the medulla part of hindbrain close to the pia surface; Astro 5114 and 5115 clusters were located along the subventricular zone, extending to the olfactory bulb and were colocalized extensively with the inhibitory immature neurons (**Figure 3d**), consistent with previous observations that the migratory steam of neuroblasts generated in the subventricular zone are ensheathed by cells of astrocytic nature^65–67, 70^. Although not all enumerated here, essentially every Astro cluster showed unique spatial distributions (**Figure 3d**). The Astro-like Bergmann cells were located in the cerebellum (**Figure 3d**), as expected. These results substantially expanded the knowledge of molecular diversity and spatial heterogeneity of astrocytes^25, 71^.

As expected, oligodendrocytes were enriched in the fiber tracts and were highly abundant throughout the brain stem, whereas the oligodendrocyte progenitor cells were evenly distributed across the whole brain (**Figure 3e**). At the cluster level, oligodendrocyte also showed regional specificity. For example, Oligo 5154 was enriched in the cortex, whereas Oligo 5162 was enriched in subcortical regions (**Figure 3e**).

We also observed region-specific distribution of the cells related to the ventricular systems. As expected, tanycytes and ependymal cells outlined the ventricles (**Figure 3f**). In the third ventricle, tanycytes resided in the ventral part whereas ependymal cells occupied the dorsal part (**Figure 3f**), consistent with previous knowledge^72, 73^. The primary residents inside the ventricles were CHOR cells, and a small fraction of VLMCs were also observed inside the ventricles (**Figure 3f**). Hypendymal cells were located in the subcommissural organ (SCO) at the dorsal third ventricle (**Figure 3f**).

Among the vascular cells, VLMCs showed region-specific distributions. Most VLMC clusters were restricted to pia, except for two distinct types: VLMC 5179 was enriched in the grey matter, and VLMC 5180 was located in the choroid plexus in the lateral and fourth ventricles (**Extended Data Figure 6a, Figure 3f**). ABCs resided near the VLMCs in the subarachnoid space (**Extended Data Figure 6b**). Other vascular cells (endothelial cells, pericytes and SMCs), which outline blood vessels, tended to be broadly distributed in the brain, as expected (**Extended Data Figure 6c**). Likewise, immune cells (microglia, BAMs, lymphoid cells, monocytes, and dendritic cells) were also scattered across the brain (**Extended Data Figure 6d**). As expected, OEGs were located at the periphery of the olfactory bulb (**Extended Data Figure 6e**).

### Molecularly defined brain regions - spatial modules

The comprehensive spatial distributions of the transcriptionally distinct cell populations allowed us to construct a map of molecularly defined brain regions. To this end, we defined for each cell a local cell-type-composition vector (see **Methods**) and clustered the cells using these vectors by a graph-based community-detection clustering algorithm^74^. The resulting clusters, which we termed “spatial modules”, defined groups of cells with similar local cell-type compositions. We identified 16 level-1 spatial modules and 127 level-2 spatial modules (see **Methods, Figure 4a, Extended Data Figure 7, Supplementary Table 3**).

**Figure 4.**
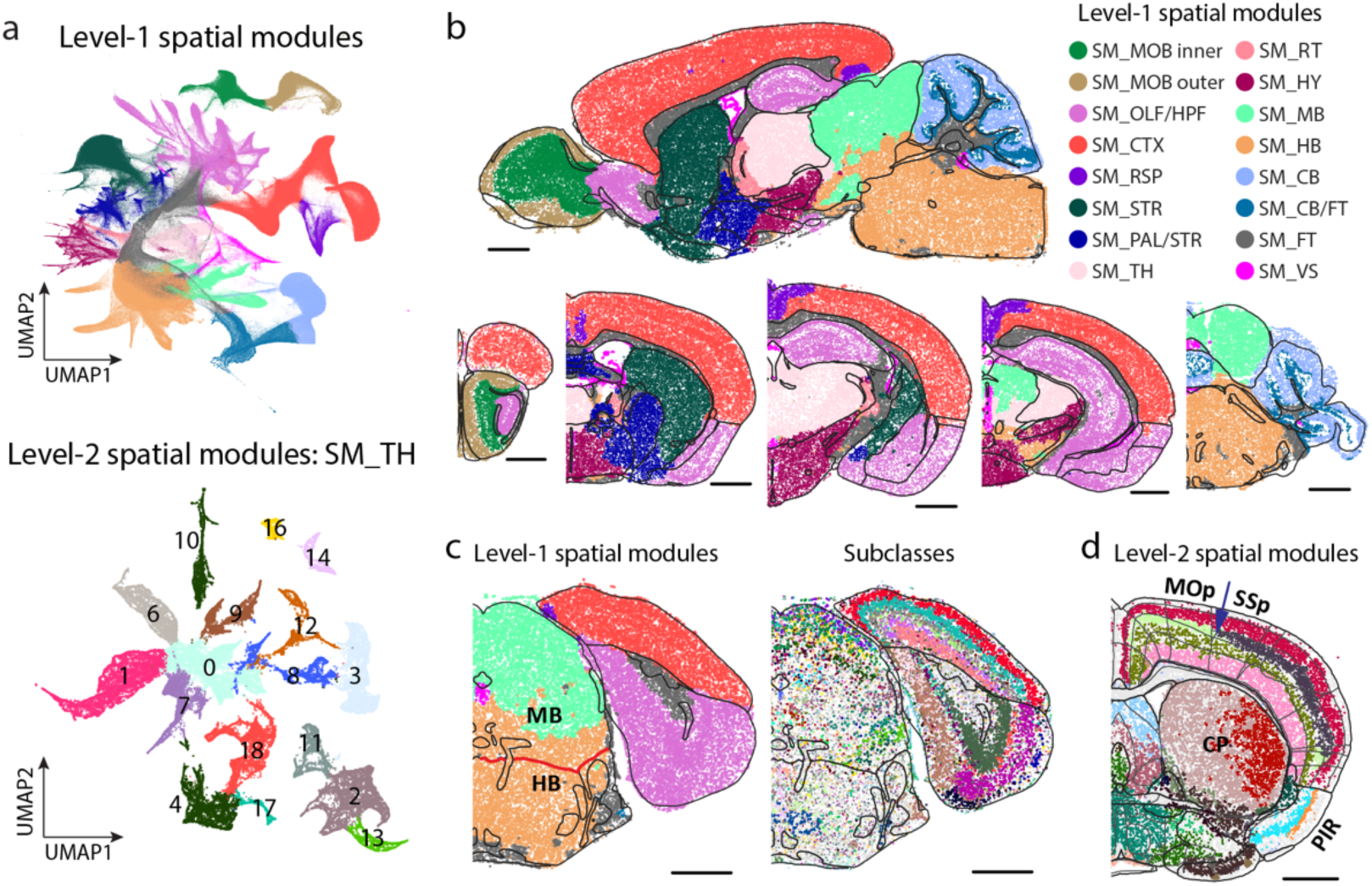
Molecularly defined brain regions. **a**, UMAP visualization of spatial modules. For any given cell, a local cell-type-composition vector is calculated, in which the elements correspond to the distance-weighted counts of cells in the neighbourhood of this cell that belong to individual cell types (see **Methods**). Clustering of cells are then performed based on their local cell-type-composition vectors to determine the spatial modules. Level-1 spatial modules are determined with the cell-type-composition determined at the subclass level; Level-2 spatial modules are then determined for each level-1 spatial module with cell-type-composition determined at both subclass and cluster levels and with only neurons considered. Top: UMAP of cells in local cell-type-composition space with cells coloured by their level-1 spatial module identities. Bottom: UMAP of cells in one of the level-1 spatial module (SM_TH, located at thalamus) with cells coloured by their level-2 spatial module identities. **b**, Spatial maps of cells, coloured by their level-1 spatial module identities shown in one sagittal and multiple coronal sections. **c**, Spatial maps of cells in one coronal section coloured by level-1 spatial module identities (left) and by cell subclass identities (right). The black lines mark the major brain region boundaries defined in the Allen CCF, and the boundary between midbrain and hindbrain defined in the CCF is highlighted in red. **d**, Spatial map of cells coloured by level-2 spatial module identities in one coronal section. The black lines mark major brain region boundaries, and the thin gray lines mark the subregion boundaries defined in the Allen CCF. The boundary between the primary motor cortex (MOp) and primary somatosensory cortex (SSp) is indicated by the blue arrow. Scale bars in **b-d**: 1 mm.

The level-1 spatial modules segmented the brain into areas that largely coincide with the major brain regions defined in the Allen CCF (**Figure 4b**). One notable discrepancy was the boundary between midbrain and hindbrain (**Figure 4c**). This discrepancy originated from the gradual changes of cell-type compositions from midbrain to hindbrain, making an unambiguous determination of midbrain-hindbrain boundary challenging. At level-2, many spatial modules were also consistent with the sub-regions defined in the Allen CCF, but we observed more discrepancies between the two at this level (**Figure 4d**). There could be multiple reasons for the discrepancies. On the one hand, our spatial module delineation was based on the cell types defined by transcriptome-wide expression profiles of individual cells and hence have a higher molecule resolution than the information used in the brain-region delineation in the CCF. For example, our analysis segmented the caudoputamen (CP) in striatum into a lateral and medial spatial module, whereas such division is not shown in the CCF brain region annotation (**Figure 4d**). Extending to the entire striatum, our analysis segmented striatum into several spatial modules, which formed a banding pattern along the dorsolateral-ventromedial axis, consistent with the banding pattern observed previously through a voxel-based clustering analysis of the Allen in situ hybridization atlas^75^. In fact, we found that further division of this region into more spatial modules were also possible, and a spatial gradient represents a more precise description of the molecular profile of this region, as described in the next section. On the other hand, we also noticed that some of the sub-region boundaries defined by connectional and/or functional information in the CCF were missing in the transcriptionally defined space modules. For example, isocortex is divided into multiple subregions in the CCF, such as the frontal cortex, primary and secondary motor cortex, primary and secondary somatosensory cortex, etc, whereas such boundaries were largely missing in the spatial module analyses except for the boundary between primary motor cortex and primary somatosensory cortex in Layer 4 (**Figure 4d**).

### Spatial gradients of molecularly defined cell types

The spatial module analysis provided a systematic characterization of molecularly defined regions in the brain. However, as a commonly encountered challenge in any clustering analysis, some of the spatial module boundaries suffered from certain level of arbitrariness, especially at locations where the cell-type composition changed gradually. Likewise, in cell-type classification, although clustering methods group cells into discrete cell types, the gene expression profiles of cells may not change abruptly across all cell-type boundaries, but rather exhibit a continuous change in some cases. Indeed, the coexistence of discrete and continuous cell-type heterogeneity has been previous observed in multiple brain regions^8, 29, 38, 76–78^, with some continuous cellular heterogeneity forming a gradient along a spatial direction^8, 29, 38, 78^.

We thus examined all cell subclasses to identify the spatial gradients of cells, in which the gene expression of cells changed gradually in space. To this end, we first systematically quantified the discreteness of clusters within each subclass (see **Methods; Figure 5a, left**). Based on this measure, most of the subclasses contained more-or-less continuously connected cell clusters, whereas subclasses with largely discrete (well separated) clusters were relatively rare (**Figure 5a, right**). In addition, among the subclasses containing largely continuously distributed cells in the gene expression space, we further identified those subclasses that exhibited a prominent spatial axis along which the gene expression profiles of cells changed gradually, indicating a spatial gradient of cells. Here we used the pseudotime^8, 79^ or the first principal component (PC1) to quantify gene expression changes. Moreover, to capture the gradients that spanned multiple subclasses, we examined subclasses that were transcriptionally similar, such as the L2/3, L4/5, L5 and L6 IT neurons, and assessed whether the gradients identified within subclasses extended into transcriptionally similar subclasses. Using this approach, we identified many spatial gradients in different brain regions. Several examples are described below.

**Figure 5.**
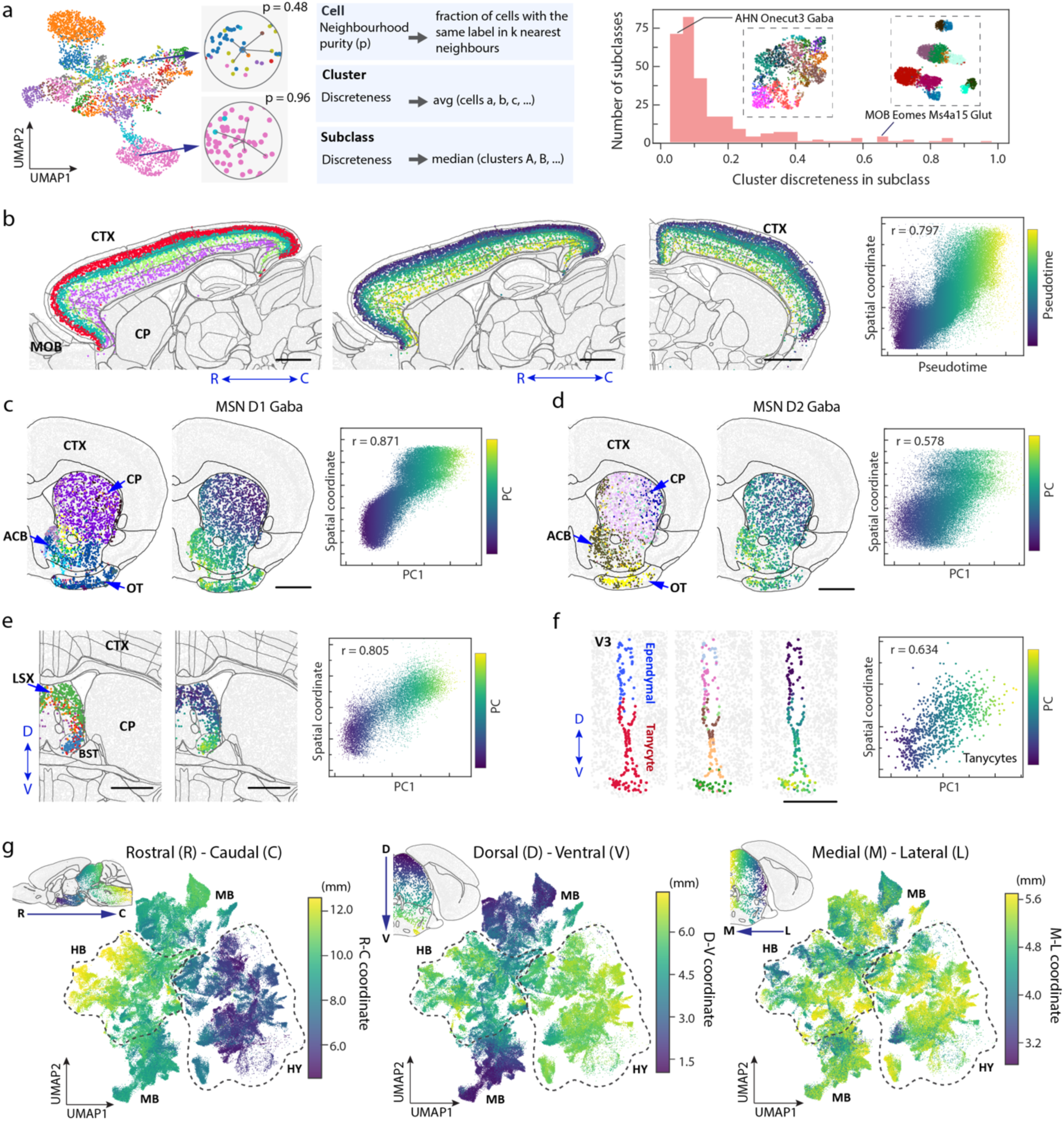
Spatial gradients of molecularly defined cell types. **a**, Left: Quantification of cluster discreteness in individual cell subclasses. For each cell in a cluster, a neighbourhood purity quantity is determined as the fraction of the cells in its neighbourhood (in the gene-expression space) that belong to this cluster. The mean neighbourhood purity quantity across all cells in a cluster is defined as the discreteness of the cluster, which gives a measure of how well separated this cluster is from the other clusters in the gene-expression space. The median discreteness of clusters is then determined for each subclass. Right: Distribution of the median cluster discreteness in individual subclasses. The UMAPs of one example subclass with high cluster discreteness (MOB Eomes Ms4a15 Glut) and one example subclass with low cluster discreteness (AHN Onecut3 Gaba) are shown. **b**, Spatial gradient of IT neurons in isocortex. From left to right: Spatial map of IT neurons coloured by subclass identities in a sagittal section; Spatial maps of IT neurons coloured by pseudotime in the same sagittal section and an additional coronal section; Correlation plot of pseudotime value versus cortical depth for individual IT neurons, coloured by pseudotime values. The Pearson correlation coefficient is r = 0.797. **c**, Spatial gradient of the D1 medium spiny neurons (MSNs) in striatum. From left to right: Spatial map of D1 MSNs coloured by subclass identities in a coronal section; Spatial map of D1 MSNs coloured by the first principal component (PC1) in the same coronal section; Correlation plot of PC1 value versus spatial coordinate for individual D1 MSNs, coloured by PC1 values. **d-f**, Same as **c** but for spatial gradients of D2 MSNs in striatum, GABAergic neurons in the lateral septal complex (LSX), and tanycytes in the third ventricle (V3). **g**, Large-scale gradient of neurons across hypothalamus (HY), midbrain (MB), and hindbrain (HB). The UMAPs are generated based on the gene-expression profiles of neurons, and individual cells are coloured by their spatial coordinates along the rostral-caudal (left), dorsal-ventral (middle), and medial-lateral (right) axes. The insets show example brain slices with cells in the regions of interest coloured by the relevant spatial coordinates. Scale bars in **b-e**: 1 mm; Scale bar in **f**: 0.5 mm.

IT neurons formed a continuous gradient across the whole isocortex, with the gene expression changed gradually along the cortical depth direction (**Figure 5b**), consistent with our previous results of IT neurons in the primary motor cortex^8^. In striatum, the D1 and D2 medium spiny neurons (MSNs) both formed a spatial gradient along the dorsolateral-ventromedial axis (**Figure 5c, d**), also consistent with previous observations^29^. In lateral septal complex (LSX), several GABAergic subclasses formed a gradient along the dorsal-ventral axis (**Figure 5e**). Similar gradients were also observed for the glutamatergic neurons in the CA1, CA3 and dente gyrus regions of hippocampus (**Extended Data Figure 8a-c**), and Tfap2d Maf Glut neurons in inferior colliculus in midbrain (**Extended Data Figure 8d**). We observed such gradients not only among neurons, but also among some non-neuronal cells. For example, tanycytes formed a continuous gradient along the dorsal-ventral axis of the third ventricle (**Figure 5f**). Overall, spatial gradients of cells were widespread in many brain regions.

We also noticed a large-scale spatial gradient spanning the hypothalamus, midbrain, and hindbrain regions. Here, we visualized the gradient in the gene-expression UMAP, where each neuron was colored by its spatial coordinate in the 3D space (**Figure 5g**). An overall rostral-caudal gradient of gene-expression change from hypothalamus to midbrain and then hindbrain, as well as a dorsal-ventral gradient from midbrain to hypothalamus and hindbrain, were apparent in these UMAPs.

### Cell-type-specific cell-cell interactions and communications

The high-resolution spatial atlas of molecularly defined cell types further allowed us to infer cell-type-specific cell-cell interactions or communications arising from soma contact, paracrine signaling, or other short-range interactions. Here, we considered cell types at the subclass level. We defined a pair of cells to be in contact or proximity if the distance between their soma centroids was within a given threshold (15 um), which was comparable to the soma size of cells in the mouse brain. We then determined, for each cell-type pair, whether the probability of soma contact or proximity observed between cells from these two cell types was statistically significantly greater than that expected from random chance. We determined the random chance (null distribution of probability) by performing local spatial-coordinate randomizations to disrupt the spatial relationship between neighbouring cells while preserving the local density of each cell type^11^ (**Figure 6a, left**). Since the stringent distance threshold may eliminate some cells that communicate through paracrine signaling, we also relaxed this distance threshold to a greater value (30 um), but for any cell-type pair identified with this relaxed distance threshold, we further required that at least one ligand-receptor pair was upregulated in the proximal cell pairs as compared to non-proximal cell pairs within this cell-type pair (**Figure 6a, right**) in order to call these cell types an interacting cell-type pair.

**Figure 6.**
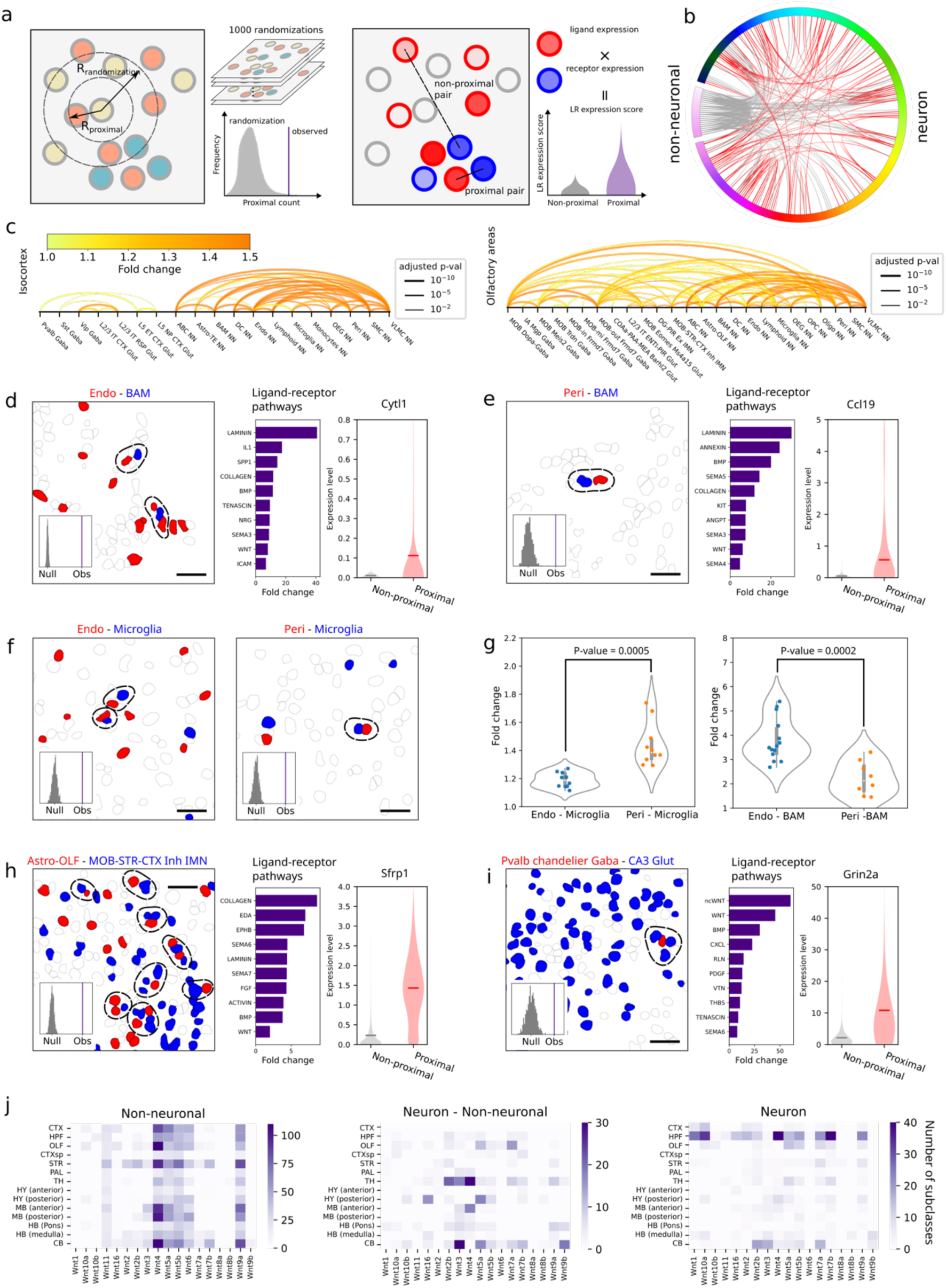
Cell-cell interactions and communications. **a**, Left: Spatial proximity analysis between a pair of cell types. A pair of cells (one from each cell type) is considered in proximity if the distance between their soma centroids is smaller than a threshold value (R_proximity_ = 15 or 30 um). The number of soma pairs that are in proximity is then determined for this cell-type pair and compared with the null distribution of this number determined by 1000 spatial-coordinate randomizations in order to determine the statistical significance of the observed soma proximity as compared to random chance. The coordinate randomization is performed locally for each cell, within R_randomization_ (chosen to be 100 um), to disrupt the spatial relationship between neighboring cells while preserving the local density of each cell type. Right: Ligand-receptor analysis. Within each cell-type pair that showed statistically significant proximity as compared to the null distribution, proximal cell pairs (soma distance < R_proximity_) and non-proximal cell pairs are identified and the distribution of the product of ligand and receptor expression levels in the proximal cell pairs are compared with that in the non-proximal cell pairs to determine the upregulation fold change and statistical significance. **b**, Predicted cell-cell interactions across the whole brain, with each line corresponding to a cell-type pair showing significant interactions by our criteria. Grey lines indicate interactions between non-neuronal cells and neurons or among non-neuronal cells; red lines indicate neuron-neuron interactions. Self-interactions are not shown. **c**, Predicted cell-cell interactions in two example brain regions, isocortex and olfactory areas. Each arc connects a cell-type pair showing significant interactions by our criteria. The colour of each arc represents the fold change between the measured number of proximal cell pairs and the mean number derived from the null distribution and the line width represents the adjusted p-value after using the Benjamini-Hochberg procedure to correct for multiple hypothesis testing. Self-interactions are not shown. Similar plots for other brain regions are shown in **Extended Data Figure 9**. **d-h**, Specific examples of predicted cell-cell interactions or communications. **d**, Interactions between endothelial cells and BAMs. Left: Example image of cells in a small area, with cells belonging to the indicated cell types shown in red and blue, and all other cells shown in grey. The proximal cell pairs are circled by dashed line. Inset shows the observed counts of the proximal cell pairs between these two cell types and the null distributions generated by local spatial randomization as described in **a**. Scale bar: 30 μm. Middle: Ligand-receptor pathways significantly upregulated in the proximal cell pair as compared to non-proximal cell pairs within the cell-type pair. A fold change of ligand-receptor expression score between the proximal cell pairs and non-proximal cell pairs is calculated for each ligand-receptor pair. When multiple ligand-receptor pairs in a pathway are upregulated, the plotted fold change value represents that of the ligand-receptor pair with the highest upregulation fold change. The pathways are rank ordered by this fold change value. Right: Expression distributions of the indicated gene in endothelial cells when they are proximal (red) or non-proximal (grey) to BAMs. **e**, Same as **d**, but for interactions between pericytes and BAMs. **f**, Interactions between endothelial cells and microglia (left) and between pericytes and microglia (right). Images and insets are as described in **d**. **g**, Fold changes of observed number of proximal cell pairs relative to the null distribution mean across different brain regions. Each data point is the fold change in a major brain region where significant interactions between the indicated cell-type pairs were observed. The p-values are calculated by two-sided Welch’s t-test. Left: Comparison between endothelial-microglia interaction and pericyte-microglia interaction. Right: Comparison between endothelia-BAM interaction and pericyte-BAM interaction. **h**, Same as **d**, but for interactions between olfactory astrocytes (Astro-OLF) and inhibitory immature neurons (MOB-STR-CTX inh IMN)). **i**, Same as **d**, but for interactions between Pvalb chandelier Gaba neurons and CA3 Glut neurons. **j**, Total numbers of unique cell types (subclasses) involved in the interacting cell-type pairs that showed upregulation of the ligand-receptor pairs involving the indicated Wnt ligands in each of the major brain regions. Left: For interactions between non-neuronal cells; Middle: For interactions between neurons and non-neuronal cells; Right: For interactions between neurons.

We applied the two approaches described above to all major brain regions and identified several hundred pairs of cell subclasses showing statistically significant interactions by our criteria, with dozens of such interacting cell-type pairs in each of the major brain regions (**Figure 6b, c**, **Extended Data Figure 9**; **Supplementary Table 4**). Most of our predicted interacting cell-type pairs contained ligand-receptor pairs, often multiple ligand-receptor pairs per cell-type pair, that showed significant expression upregulation in the proximal cell pairs as compared to non-proximal cell pairs within the same cell-type pair (**Supplementary Table 5**).

Our predicted cell-cell interactions included interactions among non-neuronal cells, between non-neuronal cells and neurons, and among neurons. Below, we describe a couple of examples in each of these three categories. As examples in the first category, we observed interactions between vascular cells and immune cells. For example, both endothelial cells and pericytes showed significant interactions with boarder-associated macrophages (BAMs) by our criteria (**Figure 6d, e**). Interestingly, in both cases, the ligand-receptor pairs that showed the most pronounced upregulation in the proximal vs. non-proximal cell pairs were in the laminin signaling pathway (**Figure 6d, e**). Laminins at the endothelial basement membrane can promote monocyte differentiation to macrophages^80^. It is thus interesting to surmise that our observed upregulation of laminin ligand-receptor pairs in these interacting cell pairs might play a role in regulating the pool of macrophages in the brain. We also observed significant interactions between microglia and these two vascular cell types (**Figure 6f**). Interestingly, compared to endothelial cells, pericytes exhibited a higher probability to interact with microglia, whereas an opposite trend was observed for the probabilities of their interactions with BAMs (**Figure 6g**).

We also observed significant interactions between neurons and non-neuronal cells. For example, astrocytes and inhibitory IMNs showed significant interactions or communications in the olfactory bulb (**Figure 6h**) and the proximal astrocyte-IMN cell pairs showed upregulated expression of ligand-receptor pairs in various pathways (**Figure 6h**). It has been shown previously that neuroblasts migrating from the subventricular zone to the olfactory bulb interact with cells of astrocytic nature along the RMS^65–67, 70^. Whether our observed interactions between IMNs and astrocytes in the olfactory bulb is related to the interactions between neuroblasts and astrocytes in the RMS remains an open question. We also observed significant interactions between astrocytes and excitatory IMNs in hippocampal formation (**Extended Data Figure 10a**). Many additional astrocyte-neuron interactions were observed across various brain regions (**Figure 6c**, **Extended Data Figure 9**). However, it is worth noting that many astrocyte-neuron interactions may also be missed by our analysis because astrocytes often interact with neurons through their processes instead of cell bodies.

Although our approach was designed to predict soma-contact-based or short-range interactions and hence is unlikely to capture long-range synaptic communications between neurons, our analysis also predicted interactions between some neuronal subclasses, for example, between Pvalb chandelier Gaba neurons and CA3 Glut neurons in hippocampal formation (**Figure 6i**) and between MSN D1 Sema5a Gaba neurons and Sst Chodl Gaba neurons in striatum (**Extended Data Figure 10b**). Interestingly, the proximal pairs of chandelier neurons and CA3 Glut neurons showed pronounced upregulation of ligand-receptor pairs in the Wnt pathways (**Figure 6i**). Wnt signaling is known to be important for hippocampal functions^81^, as well as dysfunction in neurological disorders, such as spatial memory impairment and anxiety-like behavior^82^. Chandelier neurons and CA3 Glut neurons have been previously implicated in these neurological disorders^83, 84^, but whether our observed interactions between chandelier and CA3 Glut neurons are involved in these disorders awaits future investigations.

Given the importance of Wnt signaling in brain development, function, and diseases^85, 86^, we performed a systematic quantification of our predicted involvement of various Wnt ligands in cell-cell interactions in different brain regions. Interacting non-neuronal cells primarily showed upregulation of *Wnt4*, *5a*, *5b*, *6*, and *9a*, much more prominently than the other Wnt ligands, across nearly all brain regions (**Figure 6j, left**). On the other hand, the usage of Wnt signaling in neuron-neuron communications and in communications between neurons and non-neuronal cells showed high regional specificity. For neuronal communications, Wnt signaling was highly enriched in hippocampal formation, in particular, involving the ligands *Wnt 1*, *4*, *7*, and *10a* (**Figure 6j, right**), and this observation corroborates the important roles of Wnt signaling in hippocampal functions such as memory formation^81^. Communications between neurons and non-neuronal cells showed enrichment of Wnt signaling in thalamus and cerebellum (**Figure 6j, middle**). For example, the *Wnt 4*, *2b* and *3* ligands were prominently used in thalamus and the *Wnt 3, 5a,* and *9b* ligands were prominently used in the cerebellum (**Figure 6j, middle**). Overall, among the ligand-receptor pairs that we observed to be upregulated in interacting cells in the brain, Wnt, laminin, collagen, semaphoring, and BMP-related pathways were among the most broadly used pathways (**Extended Data Figure 10c**).

In addition to ligands and receptors, we also identified other genes that were upregulated in the predicted interacting cell pairs. In each pair of interacting cell types, the proximal cell pairs often exhibited upregulation of many genes compared to the non-proximal cell pairs (**Supplementary Table 6**). Below, we illustrate this with one or two examples for each of the three major categories of interactions, non-neuronal – non-neuronal, neuronal – non-neuronal, and neuronal – neuronal interactions. For example, some cytokines were upregulated in vascular cells proximal to BAMs (e.g. *Cytl1* in endothelial cells and *Ccl19* in pericytes) (**Figure 6d, e**). These cytokines are known to be chemoattractants for macrophages^87, 88^. Our observations thus suggest the possibility that vascular cells in the brain may use these cytokines to recruit macrophages. As another example in the first category, genes involved in elastic fiber assembly, including *Eln*, *Fbln2*, and *Fbln5*, were significantly upregulated in endothelial cells proximal to SMCs (**Extended Data Figure 10d**), consistent with previous findings that endothelial cells make elastic fibers that inhibit the growth of SMCs^89^. We further observed that *Pi16* was also significantly upregulated in endothelial cells proximal to SMCs (**Extended Data Figure 10d**). Although the function of *Pi16* in this interaction is unknown, *Pi16* has been shown to inhibit the growth of cardiomyocytes^90^, a muscle cell type in the cardiovascular system. We thus hypothesize that *Pi16* expressed by endothelial cells may be a growth inhibitor of SMCs, possibly working in conjunction with the genes involved in elastic fibers. As an example in the second category – interactions between neurons and non-neuronal cells, we observed that *Sfrp1*, a Wnt signaling modulator^91^, was upregulated in astrocytes proximal to inhibitory IMNs in the olfactory bulb (**Figure 6h**). A recent study showed that *Sfrp1* expressed in OPCs in the human brain can inhibit the proliferation of neural stem cells^92^. Our results suggest the possibility that astrocytes may use *Sfrp1* to modulate Wnt signaling and regulate adult neurogenesis. Finally, as an example in the neuronal interaction category, we observed that the glutamate receptor Grin2a was upregulated in Pvalb chandelier neurons proximal to CA3 Glut neurons (**Figure 6i**). Wnt signaling is known to be important for maintaining synaptic functions in the adult brain^93^. Our observations of Wnt ligand-receptor upregulation in the proximal chandelier - CA3 Glut neuron pairs and Grin2a upregulation in chandelier cells proximal to CA3 Glut neurons (**Figure 6i**) suggest the possibility that communications between these neurons may affect the synaptic properties of chandelier neurons by upregulating the Grin2a gene. Although we discussed here only one or a few example genes, often many genes were upregulated in each predicted interacting cell-type pair, providing a rich resource for generating hypotheses of the functional implications of these cell-cell interactions.

## Discussion

In this work, we generated a spatial atlas of molecularly defined cell types across the whole mouse brain with high molecular and spatial resolution. By imaging ∼8 million cells with MERFISH and integrating the MERFISH data with a scRNA-seq dataset containing ∼4 million cells, we determined the spatial organization of >5,000 transcriptionally distinct cell clusters, which were grouped into ∼300 cell subclasses. Registration of the MERFISH images to the Allen mouse brain CCF allowed us to place the imaged cells in a common coordinate framework with each cell containing high-dimensional information, including spatial coordinates, cell-type identity, and transcriptome-wide gene expression profile (>1,100 genes measured by MERFISH and other genes imputed). This CCF registration further allowed us to determine the composition and spatial organization of transcriptionally distinct cell types in each individual brain region defined in the CCF. Analysis of the spatial relationship between cell types and correlated gene expression between proximal cells further allowed us to infer hundreds of cell-cell interactions or communications, as well as the potentially molecular basis and functional implications of these interactions.

This whole-brain cell atlas provides a comprehensive reference of the molecular diversity and spatial organization of cells in the mouse brain. Our results highlight an extraordinary diversity of neurons, comprising >5,000 transcriptionally distinct neuronal cell clusters belonging to 283 subclasses, which is accompanied by a similarly high level of spatial heterogeneity. Most of the molecularly distinct neuronal cell types exhibit distinct spatial distributions. At the subclass level, individual cell types exhibit strong enrichment, if not locate exclusively, within one of the 11 major brain regions. In the cases when a subclass of cells spans multiple brain regions, these regions are often spatially connected. At the finer scale, transcriptionally distinct neuronal clusters within individual subclasses also tend to adopt different spatial distributions from each other. We also observed different level of diversity and distinct spatial organization in different brain regions. Overall, the telencephalic regions (olfactory areas, isocortex, hippocampal formation, cortical subplate, striatum, and pallidum) show lower diversity of cells in each region, whereas the hypothalamus, midbrain and hindbrain exhibit higher cellular diversity with each region containing a substantially higher number of transcriptionally distinct cell types. This is not simply because these regions are made of a greater number of sub-regions with a simple cell-type composition. The neighbourhood of each cell also shows a substantially higher level of local cell-type complexity in these regions than in the telencephalic regions. Moreover, cells in these regions exhibit complex spatial organization with transcriptionally distinct cell types often assume irregularly shaped, partially overlapping spatial distributions. On the other hand, spatial organization of cells shows a higher level of regularity in the telencephalic regions, such as the layer-specific distribution of cortical neurons.

The spatial distributions of the transcriptionally distinct neuronal cell types allowed us to divide the brain into molecularly defined brain regions, which we termed spatial modules. Our spatial module delineation shows both similarities and differences to the brain regions defined in the current Allen CCF. The differences are in part because of the higher molecular resolution in our spatial module analysis, which provides a high-resolution refinement to the CCF region annotation in some brain areas. However, we also note that some functionally or connectionally defined brain-region segmentation shown in the CCF are missing in our spatial module delineation. In the meantime, we also observed many spatial gradients in the brain where the cell-type composition and molecular profiles of cells change gradually in space. Such spatial gradients can be found in many brain regions. Many of these gradients span multiple subregions within a major brain region, suggesting that some of the subregion divisions in the CCF, as well as some of the boundaries defined by our spatial-module analysis, may represent somewhat arbitrary divisions on continuous gradients. Interestingly, we also observed a large-scale gradient spanning the hypothalamus, midbrain, and hindbrain regions where the gene expression changes gradually along the rostral-caudal and dorsal-ventral axes.

Our data also provide a systematic molecular and spatial characterization of the non-neuronal cells. Non-neuronal cells account for more than half of the cells in the adult mouse brain, and this fraction varies substantially from region to region. We observed a remarkably high diversity of non-neuronal cells, comprising ∼100 transcriptionally distinct clusters belonging to 23 subclasses. It is possible that the observed diversity of non-neuronal cells is still an underestimation. For example, the whole-brain scRNA-seq data classified all microglia into a single cluster, whereas multiple different microglial states have been identified previously^94–97^. Although these additional states are often related to development, aging, and inflammation, a small population of cells in some of these states have been observed in healthy adult mice^94, 95, 97^.

Notably, many non-neuronal cell types also exhibited a highly level of regional specificity. This spatial heterogeneity is particularly pronounced for astrocytes, with the 24 astrocyte clusters each adopting a unique spatial distribution. While such regional-specific molecular profiles of astrocytes likely have a developmental origin, it is possible that the interactions of astrocytes with distinct types of neurons in different brain regions also contribute to the molecular diversity of astrocytes. An interest question arises as to whether the different molecular properties of distinct astrocytic subtypes play an important role in their function to support and modulate the activity of diverse neuronal cell types.

Our high-resolution cell atlas further enabled a brain-wide investigation of cell-type-specific cell-cell interactions or communications. Across the whole brain, we predicted interactions or communications between several hundred pairs of cell types at the subclass level. Furthermore, we identified multiple ligand-receptor pairs, as well as many other genes, upregulated in proximal cell pairs within each of these cell-type pairs. The identified ligand-receptor pairs provide potential molecular basis underlying the cell-cell interactions and the upregulated genes further suggest potential functional roles of these cell-cell interactions. These analyses thus generated a rich set of hypotheses on cell-cell communications that await validation by future experiments. It should be noted that our spatial-proximity-based analysis are designed to predict soma interaction, paracrine signaling, and other short-range interactions, and hence is unlikely to uncover long-range synaptic communications between neurons. Indeed, most of the predicted interactions from our analyses are between non-neuronal cells and neurons or among non-neuronal cells, although we also observed interactions between some neuronal cell types.

The spatial information in our MERFISH data offers unique advantages in predicting cell-cell interactions or communications. Previous large-scale predictions of cell-cell interactions or communications have been based on co-expression of ligand-receptor pairs derived from sequencing data^98^, which are prone to false positives^99, 100^. Indeed, to mitigate this problem, such predictions have often relied on validations by imaging experiment to probe whether the cells co-expressing the ligand-receptor pairs are indeed in contact or proximity. Our data inherently provides such spatial information in combination with the gene expression information, and hence allows cell-cell interaction predictions with both spatial and molecular analyses, which should help reduce false positives. Nonetheless, a few factors could still cause false positives and false negatives in our analyses. On the false positive side, although we applied local position randomizations of cells to generate null distributions in order to reduce the confounding effect of colocalization of cell types in a brain structure without interactions, and we further imposed the requirement of ligand-receptor upregulation in proximal cell pairs in interaction calling, it is impossible to completely eliminate such confounding effect especially when colocalization occurs within a relatively small brain structure. Decreasing the cell-proximity distance threshold and randomization distance range could help reduce such false positives, but could also remove *bona fide* interactions in the meantime because paracrine signaling may occur over a larger distance. Our requirement of ligand-receptor upregulation in proximal cell pair, as compared to non-proximal cell pairs, for cell-cell interaction calling could also cause false negatives, because the ligand-receptor pairs mediating interactions between two cell types may be expressed at a constant level regardless of whether the cells are in proximity of each other. Interested readers could use our cell atlas as a resource and adjust the parameters and requirements in our cell-cell interaction analysis to generate a more stringent or a more inclusive list of hypotheses. It is important to note that, regardless of the parameter choice, additional experiments are needed to validate these cell-cell interaction hypotheses.

As another cautionary note, although our CCF registration of the MERFISH-derived cell atlas allows characterization of cell-type composition and organization in different brain regions. alignment errors inevitably exist in CCF registration due to the differences between individual mouse brains and the average template represented by the Allen CCFv3, as well as the deformation of tissue sections that are not completely corrected for during image alignment. Improvement in CCF-registration accuracy is an active research topic and the CCF reference itself is also actively evolving. Thus, our current CCF registration provides a starting point and future method development in this area will help improve the accuracy of CCF registration. Our high-resolution cell atlas could also serve as a resource for method development in this area.

Overall, our data provides a comprehensive, molecularly defined, and spatially resolved cell atlas of the adult mouse brain, featuring the expression profiles and spatial distributions of thousands of transcriptionally distinct cell clusters belonging to hundreds of major cell types. This reference cell atlas provides a basis for future functional studies of these distinct cell populations. Both the molecular signatures and the spatial information in the atlas provide important handles for functional interrogation of specific neuronal cell types through transgenic targeting tools and optogenetic manipulations. In addition, the predicted interactions between non-neuronal cells and neurons and among non-neuronal cells, as well as the observed ligand-receptor pairs and other genes upregulated in the interacting cell pairs, further provide hypotheses and entry points for testing the functional roles of the diverse non-neuronal cell types through genetic perturbations. Furthermore, combination of transcriptomic imaging with neuronal activity imaging under various behavior paradigms, as demonstrated previously in a few brain regions^4, 5, 101, 102^, can also help reveal the functional roles of neurons. We envision exciting future studies combining spatially resolved transcriptomic analysis with measurements of various other properties, such as epigenomic profiles, morphology, connectivity, and function of cells, as well as with systematic gene perturbation methods, to connect our understanding of the brain’s molecular and cellular architecture with its function and dysfunction in health and diseases.

## Data availability

All raw and processed MERFISH data, as well as the MERFISH codebook and probes used in this work, can be accessed via the Brain Image Library (BIL): https://doi.org/10.35077/act-bag.

## Code availability

Code for MERFISH image analysis is available at https://github.com/ZhuangLab/MERlin.

## Supplementary Information

is linked to the online version of the paper.

## Acknowledgments

We thank other Allen Institute team members for their contributions in generating the scRNA-seq data and transcriptomic cell-type taxonomy used as a reference in this study. We also thank members of the Zhuang lab and the Allen Institute for helpful discussions. This work was supported in part by the National Institutes of Health (BRAIN Initiative Cell Census Network (BICCN) Grant U19MH114830 to H.Z. and X.Z.). X.P. is a HHMI Jane Coffin Childs postdoctoral fellow. X.Z. is a Howard Hughes Medical Institute investigator.

## Author Contributions

X.Z. conceived the project. M.Z., X.P., W.J., S.W.E., Z.Y., H.Z. and X.Z. designed the experiments. M.Z., X.P., W.J., A.H., and X.Z. designed the data analyses. Z.Y. designed the MEFISH gene panel with input from M.Z., X.P., W.J., H.Z. and X.Z.. M.Z., W.J., S.W.E., and L.C. performed MERFISH experiments and image decoding. M.Z., X.P., W.J., A.H. and Z.L. performed data analysis. K.A.S., B.T., Z.Y., and H.Z. provided the scRNA-seq data and transcriptomic cell-type taxonomy. M.Z. X.P. and X.Z. wrote the paper with inputs from W.J., A.H., S.W.E., Z.L., L.C., K.A.S., B.T., Z.Y., and H.Z..

## Competing Interests

X.Z. is a co-founder and consultant of Vizgen.

## Author Information

Correspondence and requests for materials should be addressed to X.Z.

## Extended Data Figure

**Extended Data Figure 1.**
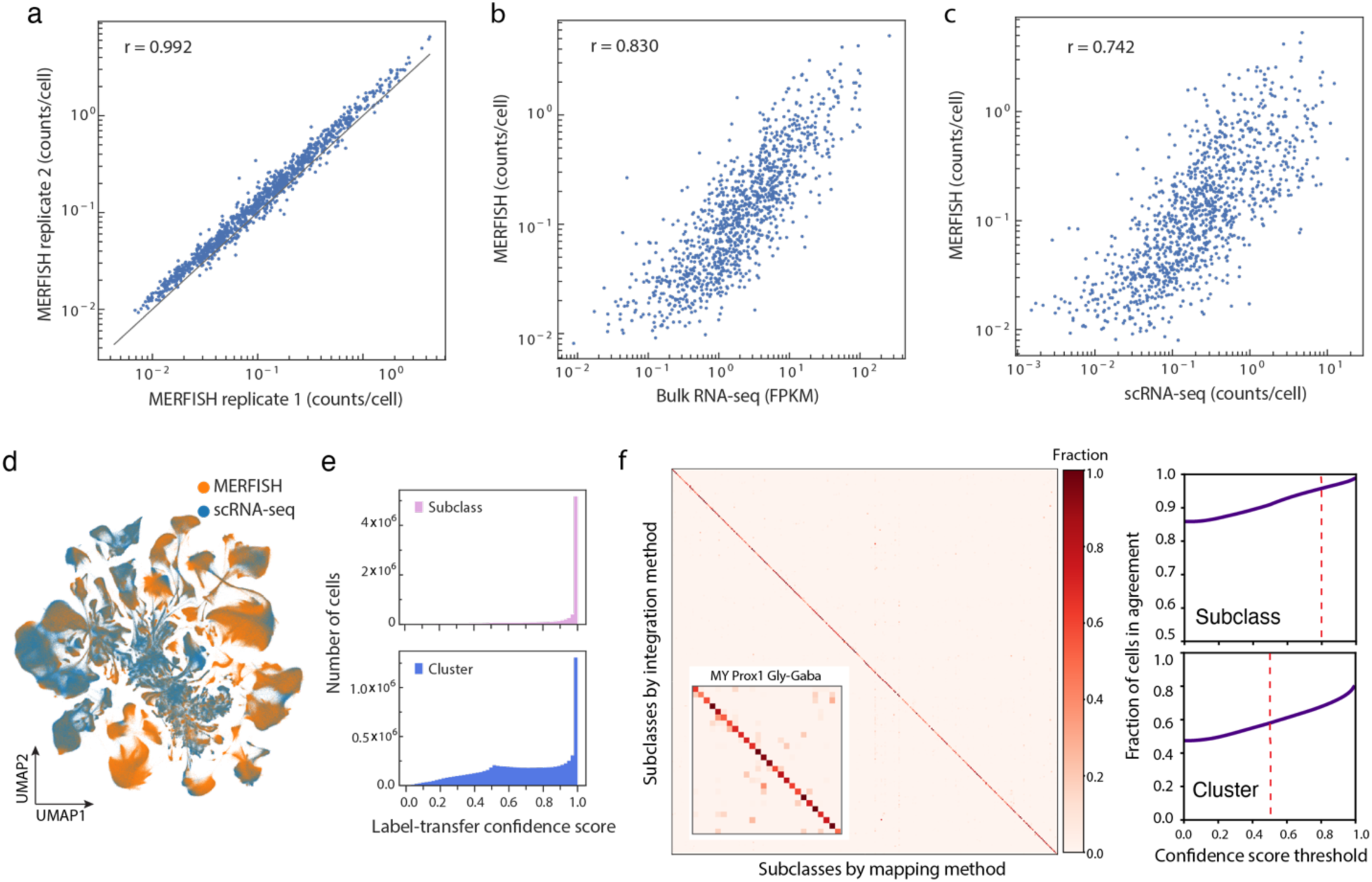
Correlation of MERFISH data and RNA-seq data and integration of MERFISH data with scRNA-seq data. **a**, Correlation plot of the average copy number per cell of individual genes measured by MERFISH from two replicate animals. The black solid line indicates equality. The Pearson correlation coefficient is r = 0.992. **b**, Correlation plot of the average copy number per cell of individual genes determined by MERFISH versus the expression levels determined by bulk RNA-seq of whole mouse brain. The Pearson correlation coefficient is r = 0.830. **c,** Correlation plot of the average copy number per cell of individual genes determined by MERFISH versus those determined by scRNA-seq of whole mouse brain. The Pearson correlation coefficient is r = 0.742. **d**, UMAP of the integrated scRNA-seq and MERFISH data with all MERFISH and scRNA-seq cells displayed. Cells are coloured by measurement modalities. **e**, Distributions of confidence scores of subclass label transfer (top) and cluster label transfer (bottom) for individual MERFISH cells. **f**, Left: Correspondence between the subclass classification of MERFISH cells determined by integration of MERFISH and scRNA-seq data (Integration method) and by identifying the scRNA-seq cluster closest to the MERFISH cells (Mapping method). Confusion matrix shows the fraction of cells from any given subclass determined by the Integration method that was assigned to individual subclasses determined by the mapping method. Insets: Correspondence plots between the cluster classification of MERFISH cells determined by the two methods for an example subclass: MY Prox1 Gly-Gaba. Right: Fraction of cells showing classification agreement between the two methods as a function of the confidence score threshold at subclass level (top) and cluster level (bottom). Red dashed lines indicate the confidence score threshold used in this work.

**Extended Data Figure 2.**
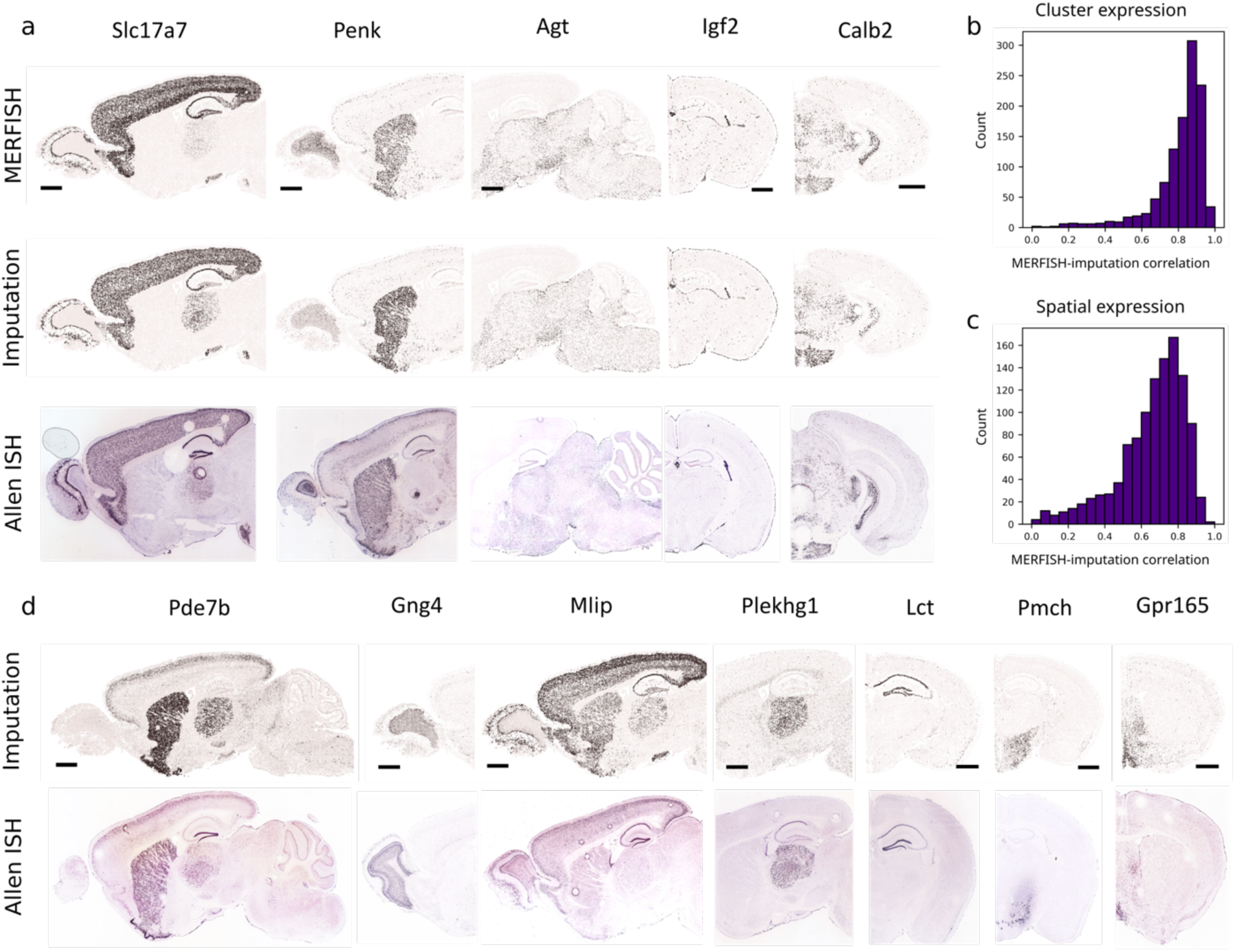
Comparison of gene-expression results imputed from MERFISH and scRNA-seq data integration with the MERFISH measurement results and Allen in situ hybridization data. **a**, Examples of spatial gene-expression patterns from MERFISH measurement (top row), imputation results (middle row), and in situ hybridization data from the Allen brain atlas (bottom row). **b,c**, The distributions of Pearson correlation coefficients between MERFISH measurement results and imputation results. **b**, For each gene, a correlation coefficient was calculated for mean expression levels in individual cell clusters between MERFISH measurement results and imputation results. **c**, For each gene, a correlation coefficient was calculated for mean expression levels of individual imaging fields of view (200 um x 200 um) between MERFISH measurement results and imputation results. Distributions over all genes in the MERFISH panel are shown. **d**, Examples of spatial gene expression patterns from imputation results (top row) and in situ hybridization data from the Allen brain atlas (bottom row). The genes in **d** were not measured by MERFISH. The Allen Brain Atlas in situ hybridization data are taken from https://mouse.brain-map.org/ (credit: Allen Institute). Scale bars in **a**, **d**: 1 mm.

**Extended Data Figure 3.**
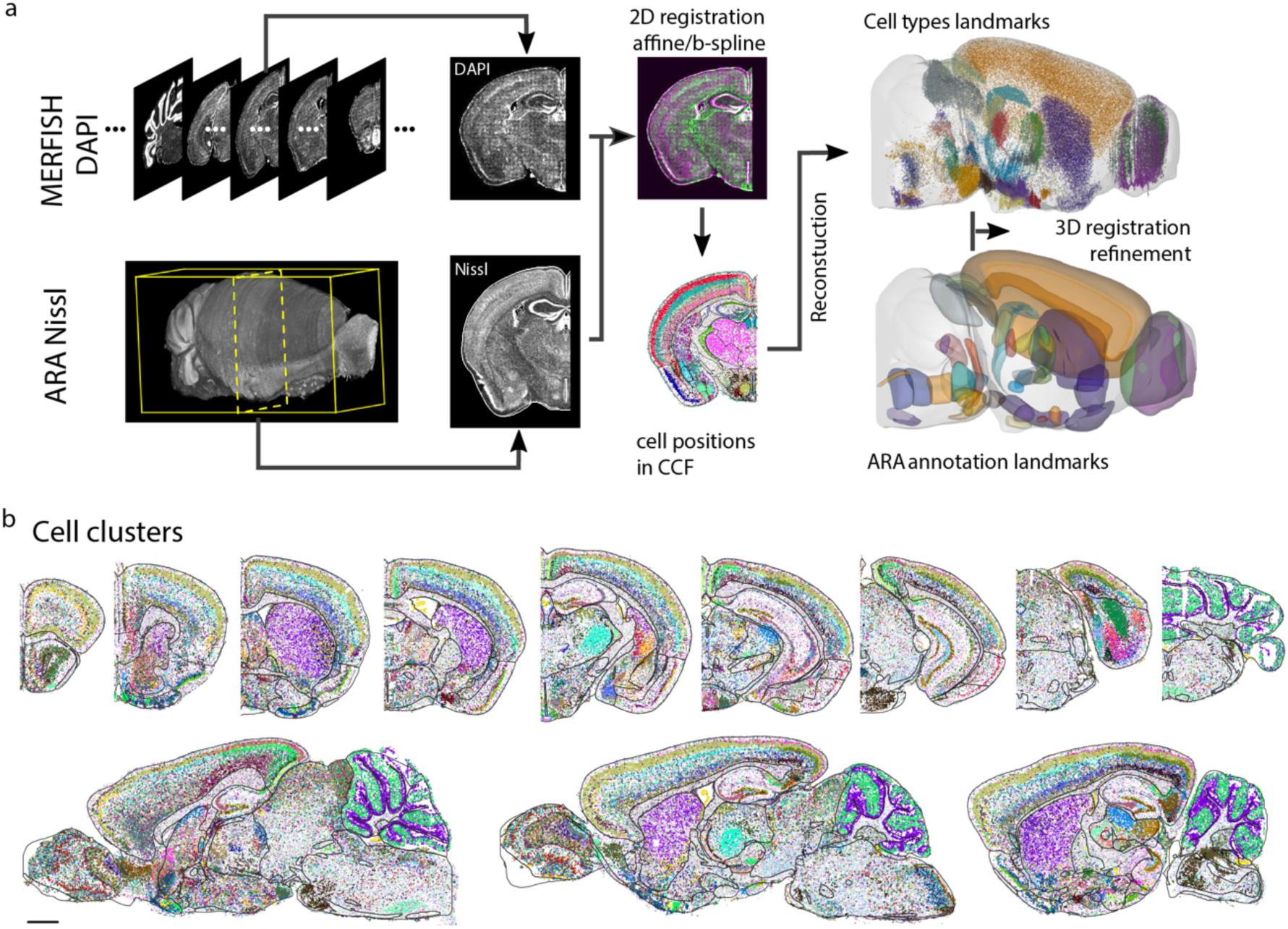
Common Coordinate Framework (CCF) registration of MERFISH images. **a**, Workflow of CCF registration of the MERFISH images. MERFISH images were registered to the Allen mouse brain CCF version 3 using a two-step procedure. First, DAPI images taken during MERFISH imaging were aligned to the Nissl template images in the Allen Reference Atlas (ARA), which allowed an initial, coarse alignment of the MERFISH images to the Allen CCF. Second, cell-type with known locations in the CCF were selected as landmarks (e.g., layer-specific cortical neurons, neurons in the dente gyrus, etc.) and used to refine the CCF alignment (see **Methods** for details). **b**, Spatial maps of cells in the same coronal and sagittal sections as shown in Figure 1c, but with cells coloured by their cluster identities. The black lines mark the major brain region boundaries defined in the CCF. Scale bar: 1 mm.

**Extended Data Figure 4.**
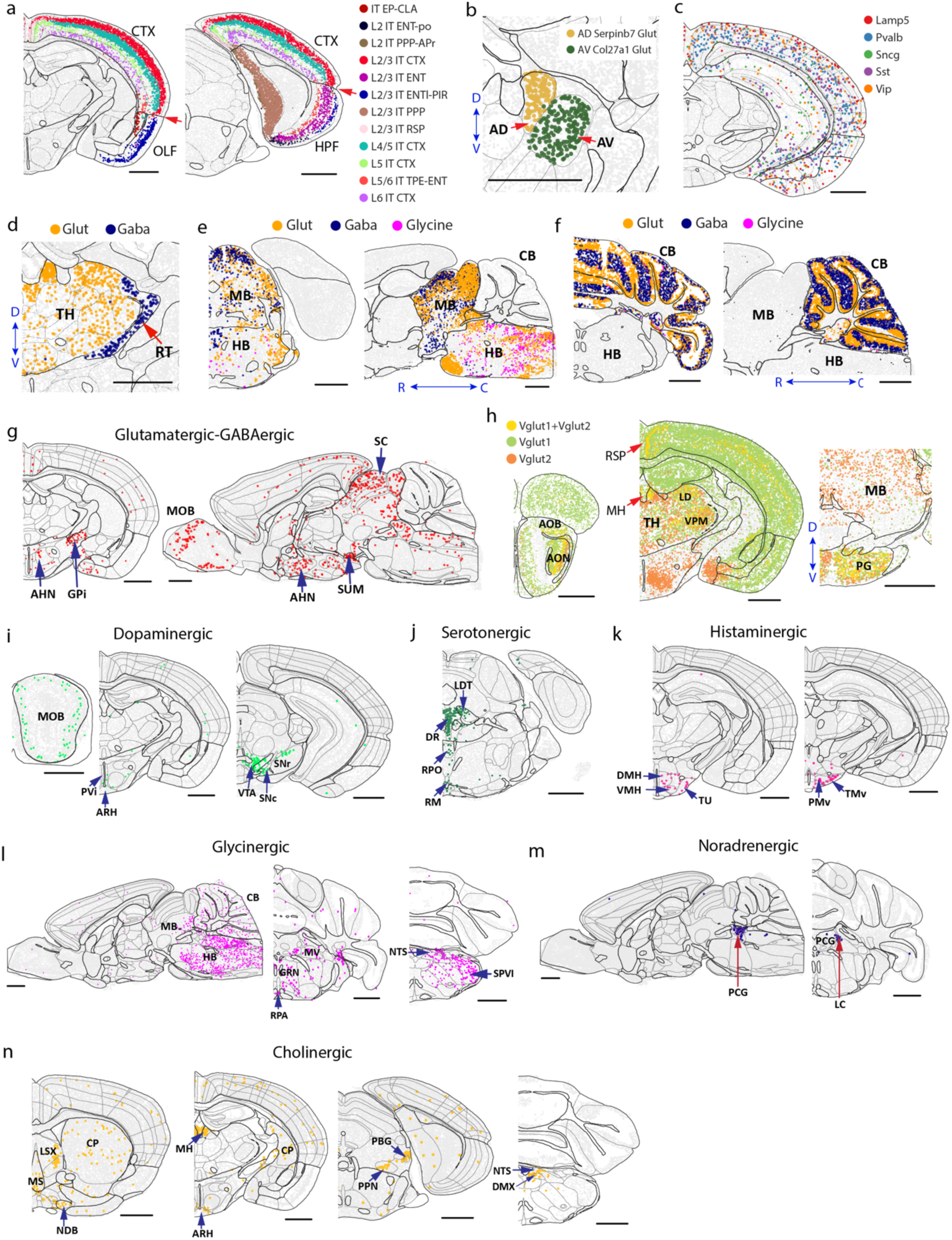
Spatial distributions of different neuronal cell types and neurotransmitter usage. **a**, Spatial distributions of different IT subclasses showing the separation between the IT neurons in isocortex (CTX) and those in olfactory areas (OLF, left) and in hippocampal formation (HPF, right). Red arrows mark the boundaries between CTX and OLF and between CTX and HPF defined in the CCF. Cells are coloured by subclass identities. **b**, Spatial distributions of the two subclasses, AD Serpinb7 Glut and AV Col27a1 Glut, in the anterodorsal (AD) and anteroventral (AV) nucleus of the thalamus, respectively. **c**, Spatial distributions of five inhibitory neuronal subclasses, marked by *Lamp5*, *Pvalb*, *Sst*, *Vip*, and *Sncg*, across CTX, HPF, OLF and cortical subplate (CTXsp). **d**, Spatial distributions of glutamatergic and GABAergic neurons in thalamus, showing GABAergic neurons in the reticular nucleus (RT) and glutamatergic neurons in the rest of thalamus. **e**, Spatial distributions of glutamatergic and GABAergic neurons, including the glycinergic neurons, in midbrain and hindbrain shown in one coronal and one sagittal section. **f**, Spatial distributions of glutamatergic and GABAergic neurons, including the glycinergic neurons, in cerebellum shown in one coronal and one sagittal section. **g,** Spatial distributions of neurons co-expressing *Vglut* (*Slc17a6*, *Slc17a7* or *Slc17a8*) and *Vgat* (*Slc31a1*) shown in one coronal and one sagittal section. **h**, Spatial distributions of neurons expressing *Vglut1* (*Slc17a7*, green) and *Vglut2* (*Slc17a6*, orange). Neurons that co-express *Vglut1* and *Vglut2* are shown in yellow and are enriched in areas such as the anterior olfactory nucleus (AON), accessory olfactory bulb (AOB), retrosplenial area (RSP), medial habenula (MH), pontine gray (PG), and multiple thalamus nuclei. **i-n**, Spatial distributions of dopaminergic (**i**), serotonergic (**j**), histaminergic (**k**), glycinergic (**l**), noradrenergic (**m**) and cholinergic (**n**) neurons shown in example coronal and sagittal sections. Scale bars in **a**-**n**: 1 mm.

**Extended Data Figure 5.**
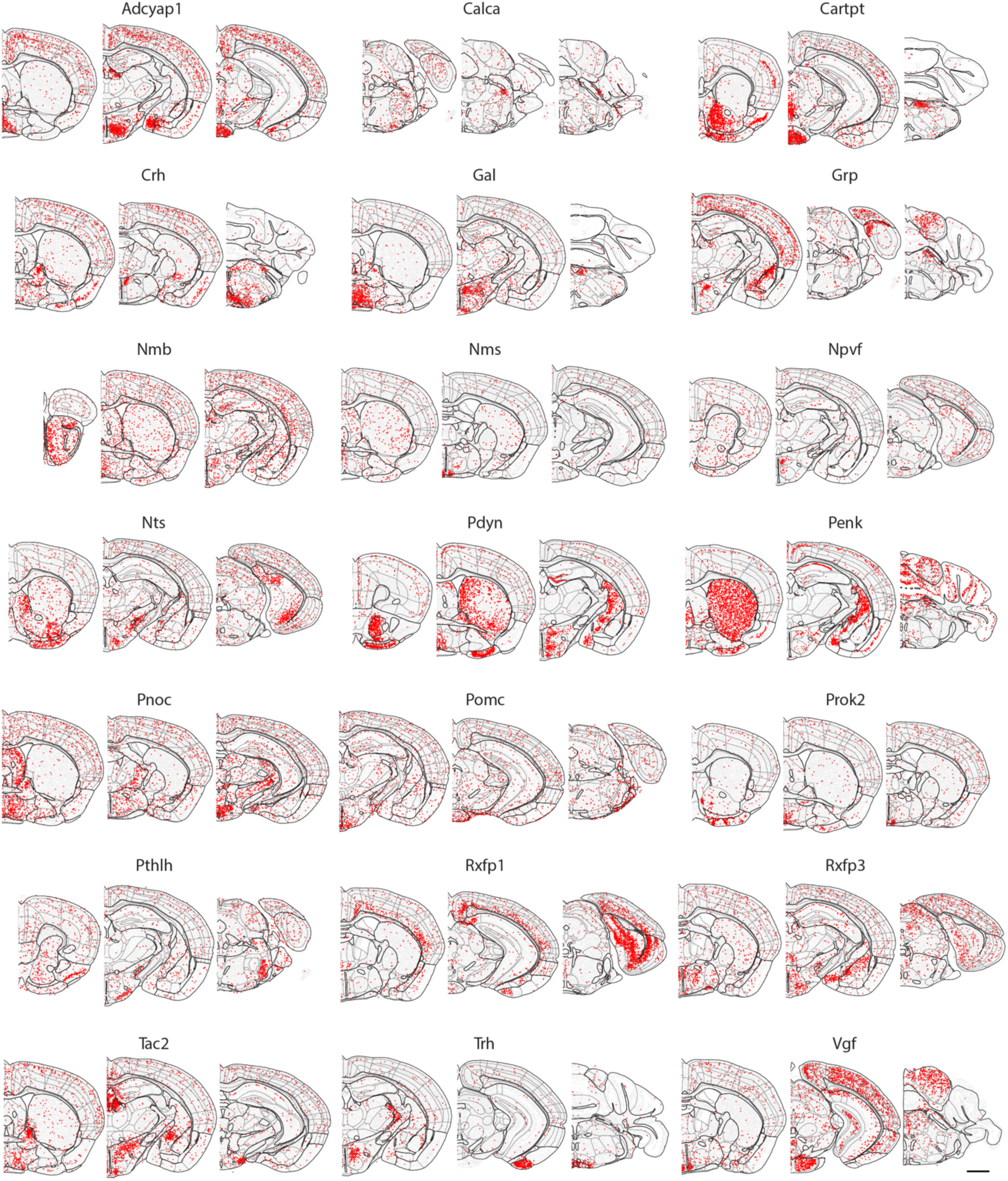
Spatial distributions of neuropeptide usage. Spatial distributions of neurons expressing various neuropeptide genes shown in multiple example coronal slices. Scale bar: 1 mm.

**Extended Data Figure 6.**
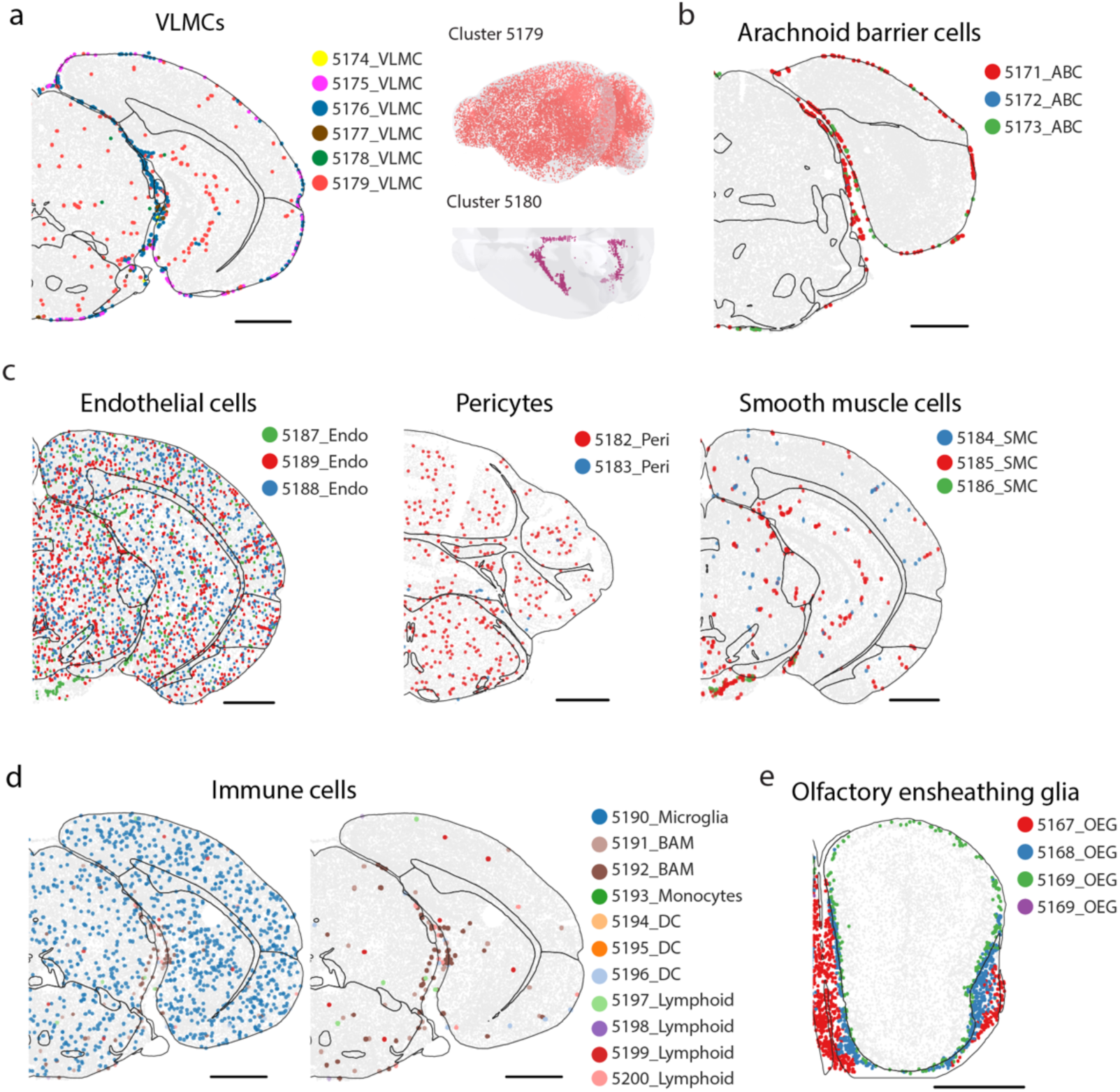
Spatial distributions of additional non-neuronal cell types. **a**, Left: Spatial distributions of VLMCs shown in one example coronal section. Right: Spatial distributions shown in the 3D CCF space for VLMC cluster 5179 (top), which is enriched in the grey matter, and cluster 5180 (bottom), which is located in the choroid plexus in the lateral and fourth ventricles. **b**, Spatial distributions of arachnoid barrier cells (ABCs) shown in one example coronal section. **c**, Spatial distributions of endothelial cells (left), pericytes (middle) and smooth muscle cells (SMCs, right), each shown in one example coronal section. **d**, Spatial distributions of immune cells shown in one example coronal section including microglia (left) and the same section but without showing microglia (right). **e,** Spatial distributions of olfactory ensheathing glia (OEG) shown in one example coronal section. Cells are coloured by cluster identities in all panels. Scale bars in **a**-**e**: 1 mm.

**Extended Data Figure 7.**
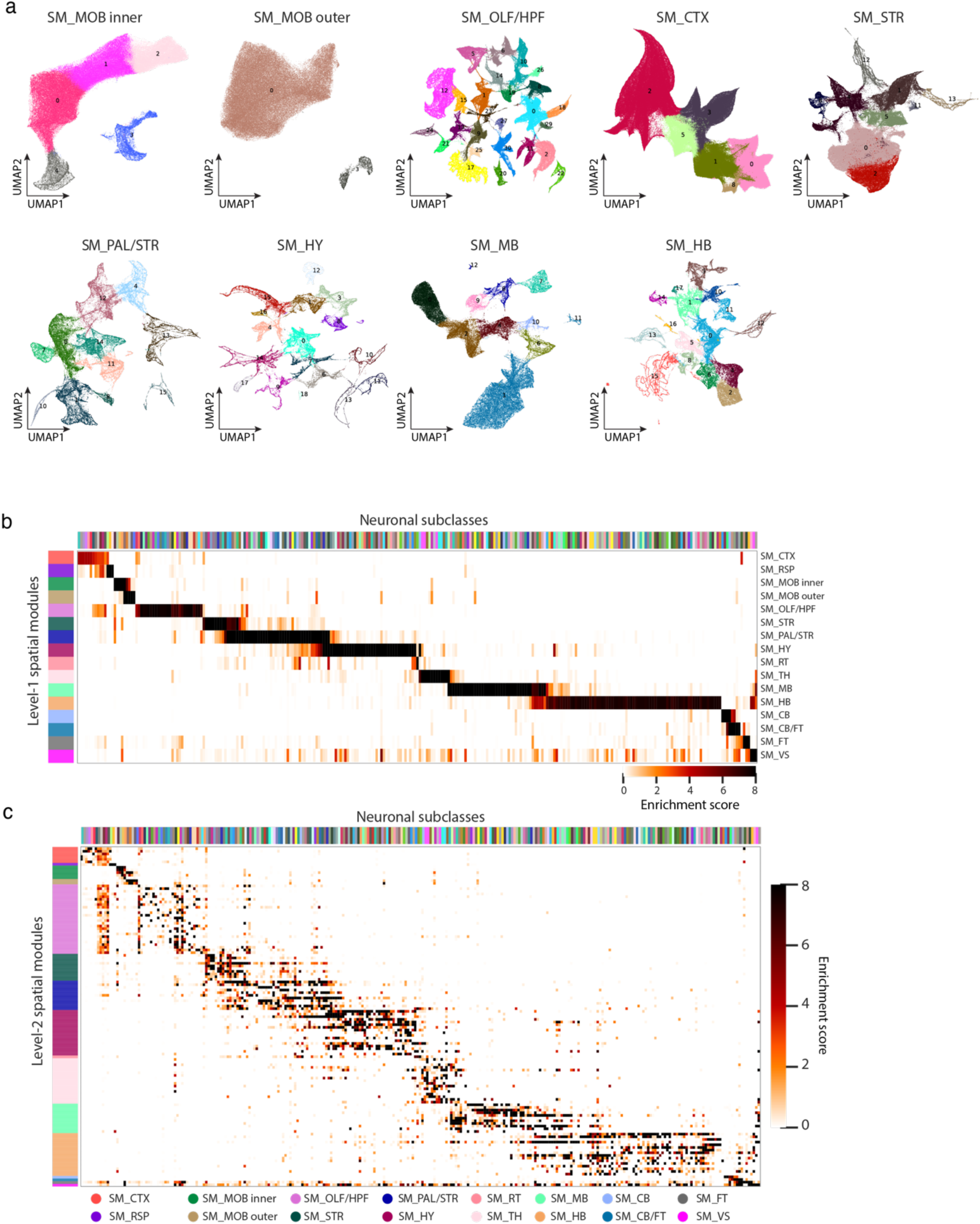
Spatial-module delineation. **a**: UMAP of cells in the other level-1 spatial module, as in Figure 4a **bottom**, with cells coloured by their level-2 spatial module identity. **b-c**, Heatmaps showing the enrichment scores of all neuronal subclasses in the 16 level-1 spatial modules (**b**) and in the 127 level-2 spatial modules (**c**). The enrichment score is defined as the fold change of the fraction of cells belong to a subclass in each individual spatial module compared to the same fraction across all spatial modules. The coloured bars at the top and on the left indicate the neuronal subclasses and spatial modul

**Extended Data Figure 8.**
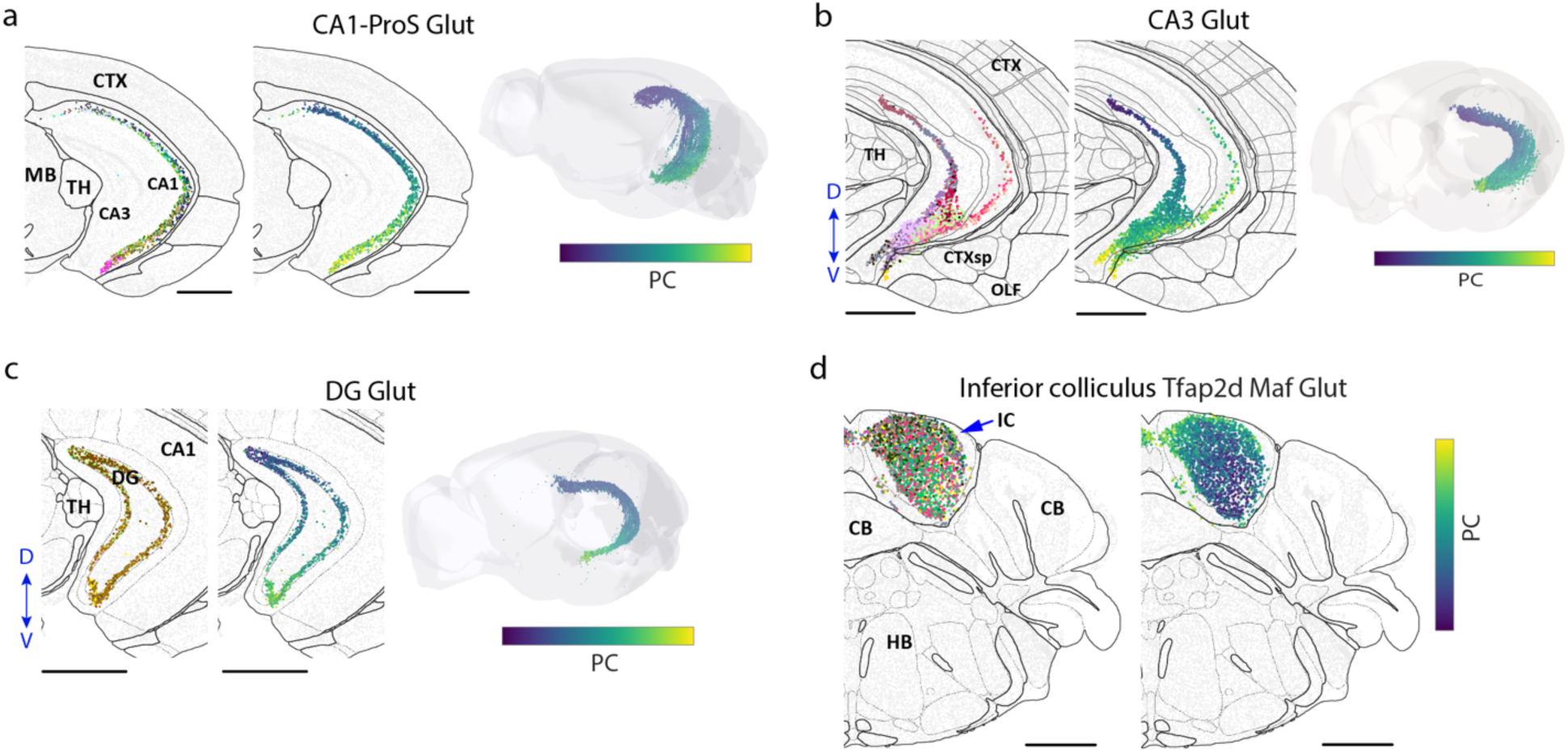
Additional examples of spatial gradients of molecularly defined cell types. **a-c**, Spatial gradients of CA3-Pros Glut neurons (**a**), CA3 Glut neurons (**b**) and DG Glut neurons (**c**) in hippocampal formation. From left to right: Spatial map of cells coloured by cluster identities in a coronal section; Spatial map of cells coloured by the first principal component (PC1) in the same section; Spatial distribution of cells colored by PC1 shown in the 3D CCF space. **d**, Spatial gradient of Tfap2d Maf Glut neurons in the inferior colliculus (IC) of the midbrain. Cells are shown in one coronal section and are coloured by cluster identities (left) and PC1 (right). Scale bars in **a**-**d**: 1 mm.

**Extended Data Figure 9.**
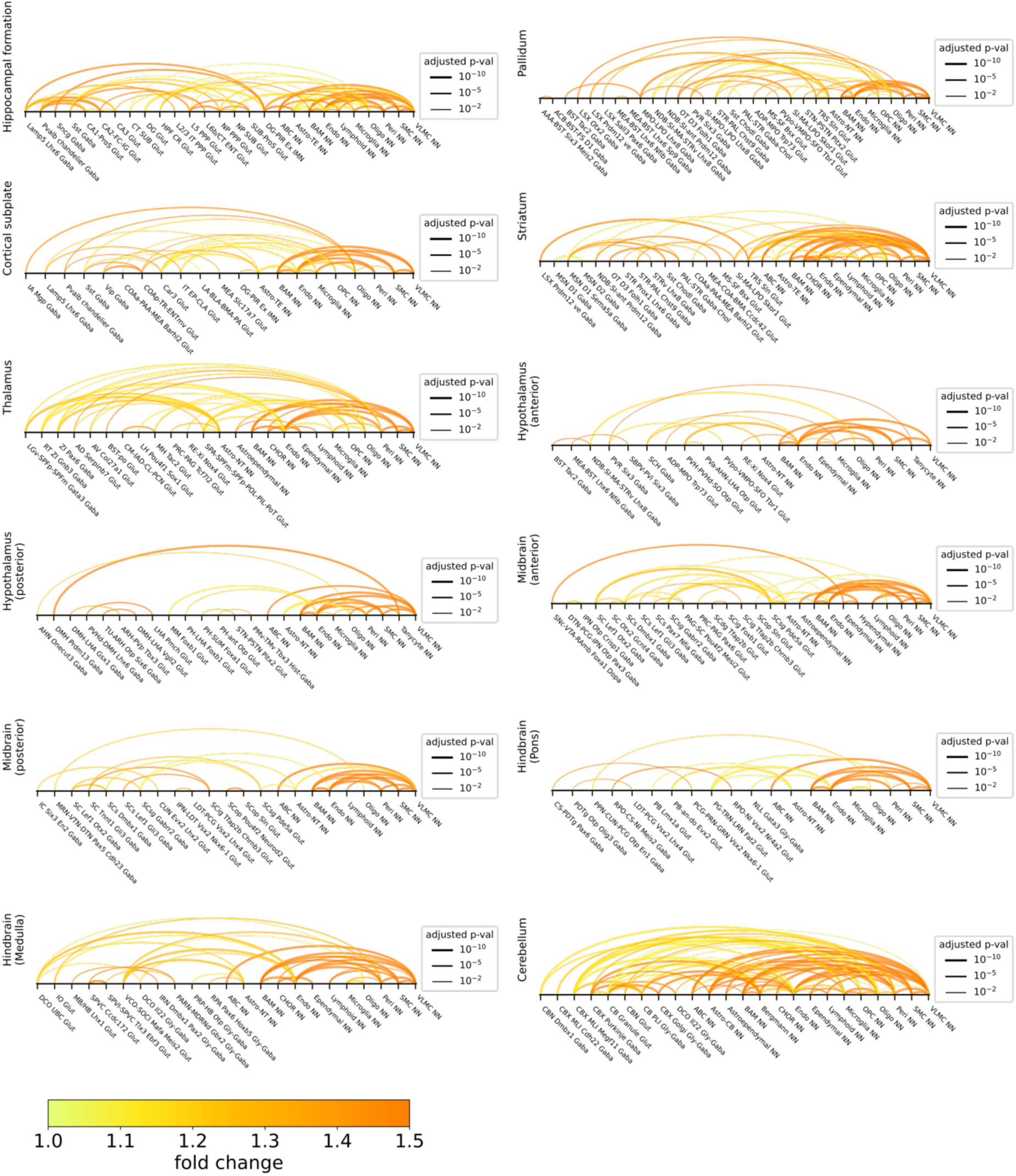
Predicted cell-cell interactions or communications in individual brain regions. Same as in Figure 6c, but for hippocampal formation, cortical subplate, striatum, pallidum, thalamus, hypothalamus (anterior and posterior parts), midbrain (anterior and posterior parts), hindbrain (pons and medulla sub-regions), and cerebellum.

**Extended Data Figure 10.**
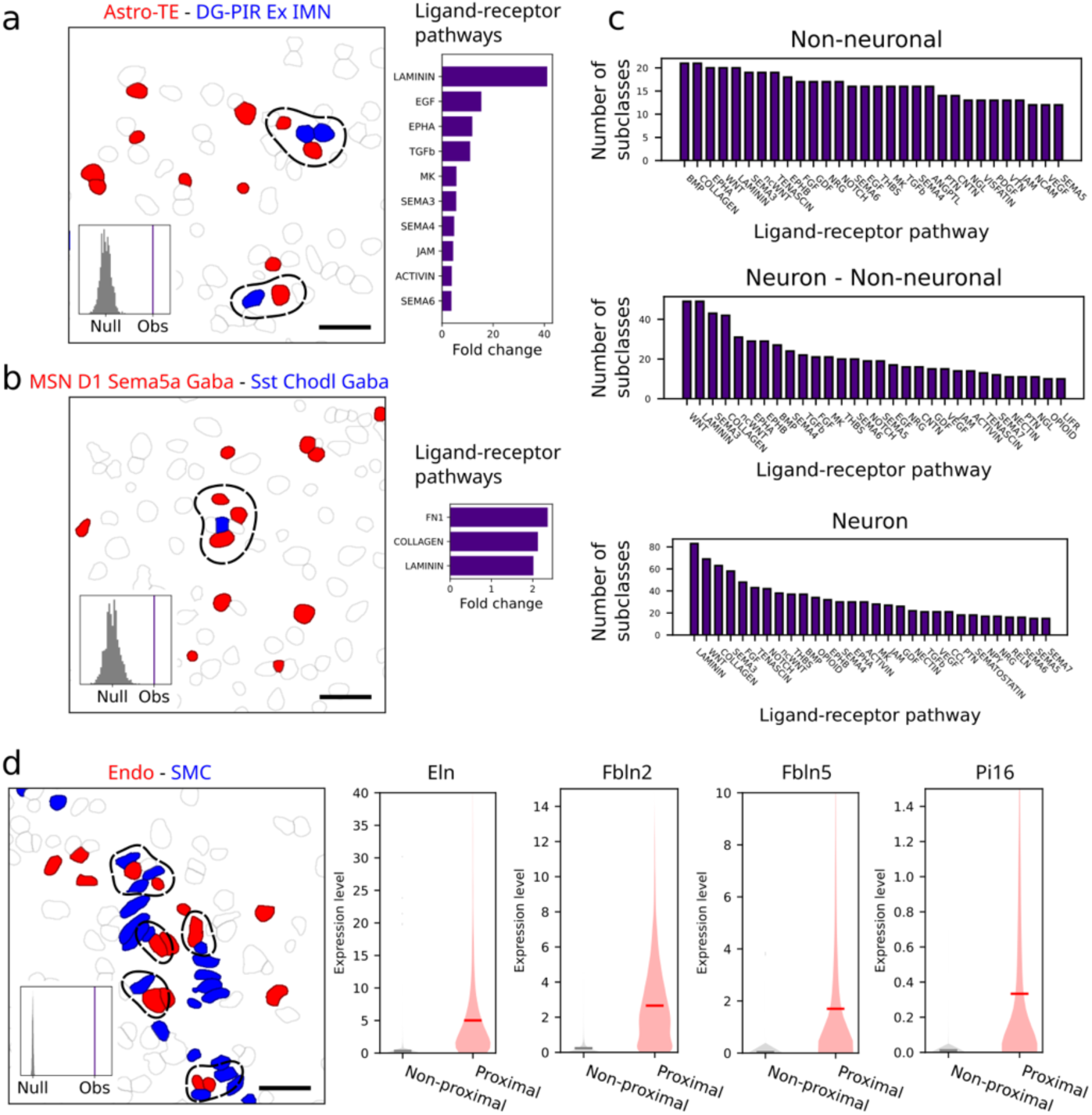
Additional examples and characterizations of predicted cell-cell interactions or communications. **a**, Interactions between astrocytes (Astro-TE) and excitatory immature neurons (DG-PIR Ex IMN). Left: Example image of cells in a small area, with cells belonging to the indicated cell types shown in red and blue and all other cells shown in grey, as described in Figure 6d. Right: Examples of upregulated ligand-receptor pathways, as described in Figure 6d. **b**, Same as **a**, but for interactions between MSN D1 Sema5a Gaba and Sst Chodl Gaba neurons. **c**, The total number of unique cell-types (subclasses) involved in the predicted interacting cell-type pairs that showed upregulation of ligand-receptor pairs in the indicated pathway across the whole brain. For each category of cell-cell interactions (interactions among non-neuronal cells, interactions between neurons and non-neuronal cells, and interactions among neurons), the top 30 ligand-receptor pathways with the highest number of cell types involved are shown. **d**, Interactions between endothelial cells and SMC cells. Left: Example image of cells in a small area, as described in Figure 6d. Right: Expression distributions of the indicated genes in endothelial cells when they are proximal or non-proximal to SMC. Scale bars in **a**, **b**, **d**: 30 μm.

## Supplementary Tables

**Supplementary Table 1 | The MERFISH genes panels imaged in this study.** Each row is a gene. “Gene” column is the gene name. “Panel” column specifies the gene panels each gene belongs to. l1: genes in the first gene panel used for animal 1; l2: genes in the second gene panel used for other animals; sequential: genes imaged in the two sequential rounds two-color FISH imaging.

**Supplementary Table 2 | Cell type compositions in major brain regions.** The “neuron” sheet contains neuronal cells, and the “non-neuronal” sheet contains non-neuronal cells. Each row is a cluster. “cluster_id” column is the cluster ID. “subclass_label” column is the subclass that the cluster belongs to. The columns after “subclass_labels” are the fractions of cells in each major brain region that belong to the indicated clusters.

**Supplementary Table 3 | Compositions of spatial modules.** The “brain_region” sheet contains brain region compositions of spatial modules. Each row is a level-1 spatial module. “spatial_modules_level_1_name” column is the name of the level-1 spatial module. The columns after “spatial_modules_level_1_name” are the fractions of cells in each spatial module that belong to individual major brain region. The “subclass” sheet contains subclass compositions of spatial modules. Each row is a level-2 spatial module. “spatial_modules_level_2_name” column is the name of the level-2 spatial module. “spatial_modules_level_1_name” column is the name of the level-1 spatial module. The columns after “spatial_modules_level_1_name” are the fractions of cells in each spatial module that belongs to individual cell subclasses.

**Supplementary Table 4 | Predicted pairs of interacting cell types.** The “15μm” sheet contains the predicted pairs of interacting cell types (subclasses) identified with the proximity distance threshold of R_proximal_ = 15μm. The “30μm” sheet contains predicted pairs of interacting cell types identified with R_proximal_ = 30μm. Each row is a predicted interacting cell-type pair in a major brain region. “subclass1” and “subclass2” columns are the labels of subclasses. “pval-adjusted” and “pval” columns are the p-values after and before the Benjamini-Hochberg multiple-hypothesis-testing correction. “z_score” column is the z-score of the measured number of proximal cell pairs compared to the null distribution. “proximal_count” column is the measured number of proximal cell pairs with soma centroid distance < R_proximal_. ”permutation_mean” column is the mean count of proximal cell pairs in the null distribution. ”permutation_std” column is the standard deviation of the count of proximal cell pairs in the null distribution. “fold_change” column is the fold change of the measured proximal cell-pair count compared to the mean of the null distribution. “region” column is the major brain region of the proximity analysis. The “supported_by_LR_analysis” column in the “30μm” sheet indicates whether a pair of proximal cell types is supported by at least one upregulated ligand-receptor pair.

**Supplementary Table 5 | Upregulated ligand-receptor pairs in the predicted interacting cell-type pairs.** Each row is a ligand-receptor pair that exhibited upregulated expression in the proximal cell pairs as compared to non-proximal cell pairs in a predicted interacting cell-type pair in a major brain region. “subclass_ligand” column is the ligand-expressing subclass. “subclass_receptor” column is the receptor-expressing subclass. “LR_pair” column is the name of the ligand-receptor pair. “N_proximal_pairs” column is the number of proximal cell pairs between the two subclasses. “mean_proximal_lr_exp” column is the mean ligand-receptor expression score in the proximal cell pairs. “exp_fraction” column is the fraction of proximal cell pairs with ligand-receptor expression scores that are greater than zero. “fold_change” column is the fold change of the mean ligand-receptor expression score in the proximal cell pairs compared to the non-proximal cell pairs. “pval” and “pval-adjusted” columns are the one-sided t-test p-values before and after the Benjamini-Hochberg multiple-hypothesis-testing correction. “region” is the major brain region of the analysis. “LR_category” column is the category of the ligand-receptor pair. “pathway_name” column is the name of the signaling pathway related to the ligand-receptor pair. “ligand” column is the gene name of the ligand. “receptor” is the gene name of the receptor. When the receptor is a complex of proteins coded by multiple genes, the gene names are separated by an underscore. “ct_pair_type” column is the type of the cell-type pair. NN-NN: non-neuronal - non-neuronal pair; NN-neuron: non-neuronal - neuron pair; neuron-neuron: neuron - neuron pair.

**Supplementary Table 6 | Upregulated genes in the predicted interacting cell-type pair.** Each row is a gene that is significantly upregulated when cells from one subclass are proximal to a cell in another subclass. “subclass_gene_exp” column is the subclass that expresses the gene. “subclass_interacting” column is the subclass that is predicted to interact with the subclass that expresses the gene. “gene” column is the name of the gene. “proximal_mean” column is the mean expression in the cells that are proximal to a cell in the interacting subclass. “control_mean” column is the mean expression in the cells that are not proximal to a cell in the interacting subclass. “fold_change” column is the fold change of the “proximal_mean” over the “control_mean.” “pval” and “pval-adjusted” columns are the one-sided t-test p-values before and after the Benjamini-Hochberg multiple-hypothesis-testing correction. “region” column is the major brain region of the analysis.

## Methods

### Animals

Adult C57BL/6 male and female mice aged 57-63 days were used in this study. Animals were maintained on a 12 hour:12 hour light/dark cycle (2pm-2am dark period) with ad libitum access to food and water. Animal care and experiments were carried out in accordance with NIH guidelines and were approved by the Harvard University Institutional Animal Care and Use Committee (IACUC).

### Bulk RNA-seq of the whole mouse brain

Estimates of the average RNA expression levels of individual genes in the mouse brain were derived from the bulk RNA-seq data of the whole mouse brain. RNA was extracted and isolated using RNAqueous Micro total RNA isolation kit (Thermo Fisher, AM1931) following manufacturer’s instructions from three different whole mouse brains aged 56-63 days. RNA quality was assessed using Agilent TapeStation and samples with an RNA integrity score >8 were kept for sequencing. RNA sequencing libraries were constructed using the Kapa mRNA HyperPrep Kits and were sequenced using the Illumina NextSeq500 platform performed by the Bauer Center Sequencing Core at Harvard University.

### Single-cell RNA sequencing data of the whole mouse brain

Single-cell RNA sequencing data were generated by the Allen Institute (See companion manuscript by Yao et al. in this BICCN package). These data are available at the Neuroscience Multi-omics Archive (https://nemoarchive.org).

### Gene selection for MERFISH

In order to discriminate transcriptionally distinct cell populations with MERFISH, we designed the gene panels based on differentially expressed (DE) gene analysis using the scRNA-seq data. Genes differentially expressed between pairs of transcriptionally distinct cell clusters from the scRNA-seq data were selected based on the following criteria: the genes had ≥2-fold change in expression between the two clusters with P-value < 0.01; they were expressed in at least 50% cells in the foreground cluster, with more than 3.3-fold enrichment, in terms of the fraction of cells expressing the gene, relative to the background cluster. Top 50 genes that satisfied the criteria and ranked by P-values in each direction for every cell-cluster pair were pooled together as the DE gene candidates for the final marker gene set. We then trimmed this DE gene pool to remove the genes that were too abundant or too short and thus were potentially challenging for MERFISH imaging experiments. Specifically, we excluded the genes that can accommodate fewer than 40 hybridization probes (MERFISH encoding probes) and thus were approximately < 500 nt in length, or were expressed at an average of 3000 counts in its highest expressing cell cluster as determined by the scRNA-seq data.

To form the MERFISH gene panel, we first included 123 subclass markers based on the scRNA-seq clustering results, and 229 genes in the gene list that included transcription factors, neuropeptides, clock genes, and GPCR/interleukin/secreted proteins related genes. We then added DE genes to the panel until there were at least 3 DE genes included for each pair of cell clusters in each direction.

Two gene panels were used in the MERFISH experiments. The first contained 1124 genes and was used for Animal 1, and the second contained 1147 genes which was used for all other animals. The two panels had 1122 genes in common. Two relatively high-abundance genes in the first gene panel were not included in the second gene panel, but 25 additional genes were added into the second gene panel, which included manually picked canonical marker genes for non-neuronal cells, as well as additional neurotransmitter related genes and neuropeptide genes.

In addition to the MERFISH gene panel, we also imaged 4 other genes (*Sst*, *Vip*, *Avp*, *Pmch*) that can accommodate fewer than 40 hybridization probes or were expressed at an average of >3000 counts in its highest expressing cell cluster. These genes were imaged in two sequential rounds of two-colour FISH imaging, following the MERFISH run that imaged the 1124-gene or 1147-gene panel.

### Design and construction of MERFISH encoding probes

Encoding probes for the MERFISH gene panels were designed as previously described^4^. We first assigned to each of the 1124 genes in first gene panel a unique binary barcode drawn from a 32-bit, Hamming-Distance-4, Hamming-Weight-4 codebook. This codebook also included 116 extra barcodes as “blank” barcodes, which were not assigned to any genes, in order to provide a measure of the false-positive rate in MERFISH measurement. For the second 1147-gene panel, the additional 25 genes were each randomly assigned a barcode from the 116 “blank” barcodes.

Each MERFISH encoding probe contained one 30-nt target sequence that could specifically bind to a target gene and two 20-nt readout sequences. We designed a total of 32 readout sequences, each corresponding to one bit of the 32-bit MERFISH code. The collection of encoding probes designed to bind each gene contained the four readout sequences corresponding to the four bits that read “1” in the barcode of that gene. Each encoding probe contained two of the four 20-nt readout sequences that encode the specific barcode assigned to the gene. To design the target sequences in the encoding probes, we identified all possible 30-nt targeting regions within each desired gene as previously described^103^. In brief, for each gene, we selected 30-nt target regions that had a GC fraction between 40% and 60%, a melting temperature (T_m_) within the range of 66-76°C, and no homology longer than 15-nt to rRNAs or tRNAs. From the set of all possible 30-nt target regions for each gene, we selected 64 target regions randomly to construct encoding probes. For the transcripts that were not long enough to accommodate 64 non-overlapping target regions, we allowed these 30-nt targeting regions to overlap by as much as 20 nucleotides to increase the number of probes. We also allowed the minimum number of probes to be included to reduce to 40, and the target regions to have a GC fraction between 30% and 70%, and a melting temperature (T_m_) within the range of 61-81°C. Among the 1147 genes, 7 genes had between 40-64 probes and the remaining genes had 64 probes.

In addition, we concatenated to each encoding probe sequence two PCR primers, the first comprising the T7 promoter, and the second being a random 20-mer designed to have no region of homology greater than 15 nucleotides with any of the encoding probe sequences designed above, as previously described^103^.

With the template encoding probe sequences designed above, we constructed the MERFISH probe set as previously described^4^. The template molecules were synthesized as a complex oligo pool (Twist Biosciences) and amplified as previously described^103^.

Encoding probes for the four genes imaged using two rounds of sequential two-colour FISH were produced in the same manner, except that 48 targeting sequences were selected for each gene if possible, and one single unique readout sequence was concatenated with targeting sequences for each gene. The four readout sequences used here, one for each gene, were different from the 32 readout sequences used for the genes imaged in the MERFISH run. These probes were purchased from Integrated DNA Technologies (IDT).

The amplified encoding probes for the MERFISH run and encoding probes for the sequential two-colour FISH rounds were mixed for tissue staining.

### Design and construction of MERFISH readout probes

We used two readout probe schemes for the 32-bit MERFISH imaging plus the two sequential rounds of FISH imaging:

1. Direct readout strategy with dye-conjugated readout probes complementary to the readout sequences, as described previously8: 36 readout probes were designed, each complementary to one of the 36 readout sequences. Each readout probes were conjugated to one of the two dye molecules (Alexa750, Cy5) via a disulfide linkage. These readout probes were synthesized and purified by Bio-synthesis, stored in Tris-EDTA (TE) buffer, pH 8 (Thermo Fisher) at a concentration of 1 μM at −20 °C.
2. Two-step readout strategy with oligonucleotide adaptors, as described previously104: i) 36 adaptor probes were designed, each consisting of a sequence complementary to one of the 36 readout sequences, concatenated by two additional common readout sequences, each for one colour channel. These adaptor probes were purchased from IDT, resuspended in TE buffer, pH 8 (Thermo Fisher) to a concentration of 1 mM and stored at −20 °C. ii) Two dye-conjugated readout probes were designed, each complementary to one common readout sequence for a colour channel, and each were conjugated to one of the two dye molecules (Alexa750, Cy5 or Alexa647) via a disulfide linkage. These readout probes were synthesized and purified by IDT, stored in TE buffer, pH 8 (Thermo Fisher) at a concentration of 100 μM at −20 °C.

### Tissue preparation for MERFISH

Mice aged 57-63 days were euthanized with CO_2_, and their brain were quickly harvested and was frozen immediately in optimal cutting temperature compound (Tissue-Tek O.C.T.; VWR, 25608-930), and stored at −80 °C until sectioning. Frozen brains were sectioned at −18 °C on a cryostat (Leica CM3050 S). Continuous set of 10-µm-thick slices were collected for imaging. For Animal 1, 10-µm-thick serial coronal sections were collected from the anterior edge to the posterior edge of the brain and every 20^th^ section was kept; for Animal 2, the brain were sectioned similarly as for animal 1, but every 10^th^ coronal section was kept; for Animal 3, 10-µm-thick serial sagittal sections were collected from the midline to the lateral edge of the brain and every 20^th^ section was kept; for Animal 4, the brain were sectioned similarly as for Animal 3, but only the sections corresponding the same medial-lateral positions as the ones that showed broken regions for Animal 3 were imaged. Each coverslip contained 2-4 coronal slices or 1-2 sagittal slices. In total, 67 slices were imaged for Animal 1, 150 slices were imaged for Animal 2, 25 slices were imaged for Animal 3, and 3 slices were imaged for Animal 4. The coverslips were prepared as described previously^4^.

Tissue slices were fixed by treating with 4% PFA in 1×PBS for 15 minutes and were washed three times with 1×PBS and stored in 70% ethanol at 4 °C for at least 18 hours to permeabilize cell membranes. The tissue slices from the same animal were sectioned at the same time and were stored in 70% ethanol at 4 °C for no longer than 2 months until all the tissue sections from the same animal were imaged.

The tissue slices were then stained with the MERFISH encoding probes. Briefly, the samples were removed from the 70% ethanol and washed with 2× saline sodium citrate (2×SSC) for three times. Then we equilibrated the samples with encoding-probe wash buffer (30% formamide in 2×SSC) for five minutes at room temperature. The wash buffer was then aspirated from the coverslip, and the coverslip was inverted onto a 50 µL droplet of probe mixture on a parafilm coated petri dish. The probe mixture comprised ∼0.5 nM of each encoding probe for the MERFISH imaging, ∼ 5 nM of each encoding probe for the two sequential rounds of two-colour FISH imaging, and 1 µM of a polyA-anchor probe (IDT) in 2×SSC with 30% v/v formamide, 0.1% wt/v yeast tRNA (Life Technologies, 15401-011) and 10% v/v dextran sulfate (Sigma, D8906). We then incubated the sample at 37 °C for 36∼48 hours. The polyA-anchor probe (/5Acryd/ TTGAGTGGATGGAGTGTAATT+TT+TT+TT+TT+TT+TT+TT+TT+TT+T, where

T+ is locked nucleic acid, and /5Acryd/ is 5’ acrydite modification) was hybridized to the polyA sequence on the polyadenylated mRNAs and allowed these RNAs to be anchored to a polyacrylamide gel as described below. After hybridization, the samples were washed in encoding-probe wash buffer for 30 minutes at 47 °C for a total of two times to remove excess encoding probes and polyA-anchor probes. All tissue samples were cleared to remove fluorescence background as previously described^4, 105^. Briefly, the samples were embedded in a thin polyacrylamide gel and were then treated with a digestion buffer of 2% v/v sodium dodecyl sulfate (SDS; ThermoFisher, AM9823), 0.5% v/v Triton X-100 (Sigma, X100), and 1% v/v proteinase K (New England Biolabs, P8107S) in 2×SSC for 36-48 hours at 37 °C. After digestion, the coverslips were washed in 2×SSC for 30 minutes for a total of four washes and then stored at 4°C in 2×SSC supplemented with 1:100 Murine RNase inhibitor (New England Biolabs, M0314S) for no longer than 2 weeks prior to imaging.

### MERFISH imaging

We used home-built imaging platforms for MERFISH imaging in this study, as described previously^106^. A commercial flow chamber (Bioptechs, FCS2) with a 0.75-mm-thick flow gasket (Bioptechs, 1907-100; DIE# F18524) was used, and imaging buffer comprising 5 mM 3,4-dihydroxybenzoic acid (Sigma, P5630), 50 µM trolox quinone, 1:500 recombinant protocatechuate 3,4-dioxygenase (rPCO; OYC Americas), 1:500 Murine RNase inhibitor, and 5 mM NaOH (to adjust pH to 8.0) in 2×SSC was used for all experiment. For sagittal slices, whole tissue slices were imaged, and for coronal slices, we imaged one hemisphere plus a narrow region near the midline in the other hemisphere. Two imaging schemes were used for the two different readout strategies:

1. For the direct readout strategy, we first stained the sample with a readout hybridization mixture containing the readout probes associated with the first round of imaging, as well as a probe complementary to the polyA-anchor probe and conjugated via a disulfide bond to the dye Alexa488 at a concentration of 3 nM for imaging total poly-adenylated mRNA. The readout hybridization mixture was composed of the readout-probe-wash buffer containing 2×SSC, 10% v/v ethylene carbonate (Sigma, E26258), and 0.1 % v/v Triton X-100, supplementing with 3 nM each of the appropriate readout probes. The sample was incubated in this mixture for 15 minutes at room temperature, and then washed in the readout-probe-wash buffer supplemented with 1 µg/mL DAPI for 10 minutes to stain nuclei within the sample. The sample was then washed briefly in 2×SSC and was ready for imaging. After the first round of imaging, the dyes were removed by flowing 2.5 mL of cleavage buffer comprising 2× SSC and 50 mM of Tris (2-carboxyethyl) phosphine (TCEP; Sigma, 646547) with 15 min incubation in the flow chamber to cleave the dyes linked to the readout probes through disulfide bond. The sample was then washed by flowing 1.5 mL 2× SSC. To perform the second round of imaging, we flowed 3.5 mL of the readout probe mixture containing the appropriate readout probes across the chamber and incubated the sample in this mixture for 15 minutes. Then the sample was then washed by 1.5 mL of readout-probe-wash buffer and then 1.5 mL of imaging buffer was introduced into the chamber.
2. For the two-step adaptor readout strategy, we first stained the sample with an adaptor probe hybridization mixture containing the adaptor probes associated with the first round of imaging. The readout hybridization mixture was composed of the readout-probe-wash buffer containing 2×SSC, 30% v/v formamide (Ambion, AM9342), supplementing with 100 nM each of the appropriate adaptor probes. The sample was incubated in this mixture for 15 minutes at room temperature, washed in the readout-probe-wash buffer, and stained with a readout hybridization mixture containing 10 nM each of the two readout probes, as well as the polyA-anchor probe (Alexa488) at a concentration of 3 nM in the readout-probe-wash buffer (2×SSC, 30% v/v formamide). The sample was incubated in this mixture for 15 minutes at room temperature, washed again, and was then washed in 2×SSC supplemented with 1 µg/mL DAPI for 10 minutes to stain nuclei. Finally, the sample was washed briefly in 2×SSC and was ready for imaging. After the first round of imaging, the dyes were removed by flowing 2.5 mL of cleavage buffer comprising 2× SSC, 30% formamide and 50 mM TCEP, supplemented with unlabeled common readout probes at 100 nM each to block unoccupied readout sequences on the adaptor probes to prevent crosstalk between rounds of hybridizations. The sample was incubated in this cleavage buffer for 15 min in the flow chamber, then washed by flowing 1.5 mL readout-probe-wash buffer. To perform the second round of imaging, we flowed 3.5 mL of the adaptor probe mixture containing the appropriate adaptor probes across the chamber and incubated the sample in this mixture for 15 minutes, washed by 1.5 mL of readout-probe-wash buffer, and flowed 3.5 mL of the readout probe mixture containing the common readout probes across the chamber and incubated the sample in this mixture for another 15 minutes. Then the sample was washed again by 1.5 mL of readout-probe-wash buffer and then 1.5 mL of imaging buffer was introduced into the chamber.

In the first round of imaging, we collected images in the 750-nm, 650-nm, 560-nm, 488-nm, and 405-nm channels to image the first two readout probes (conjugated to Alexa750 and Cy5/Alexa647, respectively), the orange fiducial beads, the total polyA-mRNA signal by the polyA-anchor readout probe (Alexa488), and the nucleus signal by DAPI (405-nm channel). The latter two channels were used for cell segmentation as described below. For the second and all following imaging rounds, we collected images in the 750-nm, 650-nm, and 560-nm channels for the 2 readout probes and fiducial beads. During each imaging round, for the fiducial beads, we took a single image at one z-position for each field of view (FOV) on the surface of the coverslip using the 560-nm illumination channel as a spatial reference to correct for slight drift of the stage position over the course of imaging rounds. For imaging readout probes in the MERFISH rounds, we imaged multiple z-positions in each FOV: For Animal 1, we collected three or six 1.5-µm-thick z-stacks; for all other animals, we collected five 1.5-µm-thick z-stacks. We repeated the hybridization, wash, imaging and cleavage for all rounds to complete the 16 rounds of imaging for 32-bit MERFISH experiments. We then performed two additional rounds of two-color FISH imaging to image the four additional genes, and these images were only acquired from one z-plane per FOV. All buffers and readout probe mixtures were loaded with a home-built, automated fluidics system composed of three, 12-port valves (IDEX, EZ1213-820-4) and a peristaltic pump (Gilson, MP3).

### MERFISH image analysis and cell segmentation

All MERFISH image analysis was performed using MERlin^107^, as described previously^106^. First, we identified the locations of the fiducial beads in each FOV in each round of imaging and used these locations to determine the X-Y drift in the stage position relative to the first round of imaging and to align images for each FOV across all imaging rounds. We then high-pass filtered the MERFISH image stacks for each FOV to remove background, deconvolved them using 10 rounds of Lucy-Richardson deconvolution to tighten RNA spots, and low-pass filtered them to account for small movements in the centroid of RNAs between imaging rounds. Individual RNA molecules imaged by MERFISH were identified by our previously published pixel-based decoding algorithm using MERlin^107^. After assigning barcodes to each pixel independently, we aggregated adjacent pixels that were assigned with the same barcodes into putative RNA molecules, and then filtered the list of putative RNA molecules to enrich for correctly identified transcripts as described previously^106^ for a gross barcode misidentification rate at 5% using MERlin^107^.

We performed cell segmentation using the DAPI and total polyA-mRNA signals and a deep learning-based cell segmentation algorithm (Cellpose)^108^, as described previously^11^. For each individual z-plane, we segmented cell nuclei with the DAPI stain images with diameter parameter of 100 pixels in the “nuclei” mode. The centroid positions of the cells were then identified in each z-plane, and the centroids within distance of 2 μm in the xy direction across different z-planes were considered to be the same cell and were connected. We also segmented cell soma by the polyA images also using Cellpose with diameter parameter of 200 pixels in a “cytoplasm” mode.

We assigned unique IDs for each segmented cell and assigned individual RNAs to segmentation boundaries of the cells based on whether or not they fell within those boundaries to obtain the cell x gene matrix, i.e., the copy number of RNAs for each gene in each cell.

For the two sequential rounds of two-colour FISH imaging, we quantified the signal from these images by summing the fluorescence intensity of all pixels that fell within the segmentation boundaries of the cells associated with the imaged z-plane and normalized the signal by the areas of the cells in the z-plane.

### Preprocessing of MERFISH data

With the cell x gene matrix obtained as described above, we preprocessed the matrix by several steps: (1) The segmentation approach we used generated a small fraction of putative cells with very small total volumes due to spurious segmentation artifacts, as well as some cells that overlapped in the z dimension and were not properly separated. We hence removed the cells that had a volume of < 50 µm^3^ or > 1500 µm^3^ for the 3-z-plane measurements, the cells that had a volume of < 80 µm^3^ or > 2500 µm^3^ for the 5-z-plane measurements, and cells that had a volume of < 100 µm^3^ or > 3000 µm^3^ for the 6-z-plane measurements. (2) To remove the differences in RNA counts due to the different soma volumes captured in the images, we normalized the RNA counts per cell by the imaged volume of each cell. (3) We corrected the mean total RNA counts per cell to a same mean value (250 in this case) for each experiment. (6) We removed the cells that had total RNA counts in top and bottom 1% quantile. (8) We removed potential doublets using Scrublet^109^ as described previously. The cells with doublet score higher than 0.25 were removed as doublets, which accounted for ∼4% of the total cell number.

### Integration of MERFISH data with scRNA-seq data

We grouped MERFISH data from all four animals for integration with scRNA-seq data. Hence, only the overlapping 1122 genes between the two MERFISH gene panels used for all four animals were included in the cell x gene matrix for integration of MERFISH and scRNA-seq data and subsequent analyses.

We used the SeuratIntegration class from the ALLCools python package^35, 50^ to integrate the MERFISH dataset and the scRNA-seq dataset. The integration works by co-embedding the two datasets in a common space and finding pairs of cells from the two datasets that are close to each other in the co-embedded space. The identified close pairs are termed anchors, which were used for transferring cell type labels and imputing gene expressions from the scRNA-seq dataset to the MERFISH dataset. We performed co-embedding of the two datasets by a canonical correlation analysis (CCA) based integration method initially developed in the Seurat R package^35, 50^. In order to integrate more than ten million cells from the two datasets while achieving a fine resolution for >5,000 transcriptionally distinct cell clusters identified in the scRNA-seq data, we performed two rounds of integration.

First, we divided the cells from both datasets into 48 integration partitions. We used the scRNA-seq dataset to define the partitions. Each integration partition was a group of subclasses that were close in the transcription space. We subset the genes in the scRNA-seq dataset to the genes measured by MERFISH. Then we preprocessed the dataset using the Scanpy pipeline^110^: normalize the total count of each cell to 1000, log1p transform the counts, and scale the transformed counts to Z-scores. We reduced the dimensionality to 100 principal-component-analysis (PCA) dimensions and calculated the 15 nearest neighbors of each cell in the PCA space. From the nearest neighbor graph, we calculated a connectivity graph of subclasses where each node was a subclass, and the weight of each edge was the number of edges in the nearest neighbor graph that connected cells from the two subclasses. Then we used the direct k-way cuts method from the METIS graph partitioning library^111^ to divide the 306 subclasses into 48 integration partitions. This method aimed to evenly distribute cells into partitions while minimizing the sum weight of cut edges.

In the first round of integration, we transferred the integration-partition labels from the cells in the scRNA-seq dataset to the cells in the MERFISH dataset. We subset the genes in the scRNA-seq dataset to the genes measured by MERFISH. Then we independently preprocessed the scRNA-seq and MERFISH datasets by the Scanpy pipeline^110^: normalize the total count of each cell to 1000, log1p transform the counts, and scale the transformed counts to Z-scores. We combined the two datasets and performed PCA to reduce the dimensionality to 100. We ran CCA to co-embed the scRNA-seq cells and MERFISH cells into a 100-dimensional space. In order to co-embed the large number of cells from the two datasets, the CCA was first performed on randomly downsampled scRNA-seq and MERFISH datasets, each containing 100,000 cells. Then the CCA coordinates of the full datasets were calculated by a linear transformation from the gene expression space to the CCA space. We found the five nearest neighbors across the two datasets in the CCA space. We defined all pairs of cells from the two datasets that were mutual nearest neighbors as integration anchors. Then we used the label_transfer function from the SeuratIntegration class to transfer the integration-partition labels from the scRNA-seq dataset to the MERFISH dataset. For each MERFISH cell, the label_transfer function calculated the probability of assigning the MERFISH cell to every integration-partition based on the 100 nearest-neighbour anchor cells from the scRNA-seq dataset in the PCA space. We set the integration-partition label of a MERFISH cell to be the one with the highest probability (i.e., the integration partition that had the highest fraction of cells in the 100 nearest-neighbour anchor cells) and defined this probability as the confidence score of the transferred partition label.

In the second round of integration, we transferred subclass and cluster labels from the scRNA-seq dataset to the MERFISH dataset. We performed this round of integration for each integration-partition separately. We subset the genes in the scRNA-seq dataset to the genes measured by MERFISH, normalized the total count of each cell to 1000, and log1p transformed the counts. We used the genes that were highly variable in each integration partition. To this end, we calculated the dispersions of all the selected genes using the highly_variable_genes function from the Scanpy package^110^. Only genes with log dispersions greater than zero were kept for integration. Using the same method for the first round of integration, we transferred the subclass and cluster labels from the scRNA-seq dataset to the MERFISH dataset and calculated the confidence scores for label transfer. Because a cell-type label is transferred correctly to a cell only when both the integration-partition label and the cell-type label within the integration partition were transferred correctly, we adjusted the confidence scores of the subclass and cluster label transfer by multiplying them with the integration-partition label-transfer confidence scores.

### Imputation of transcriptome-wide gene expressions of individual cells in MERFISH images

Based on the integration of MERFISH and scRNA-seq data, we also imputed the transcriptome-wide gene expression for each cell in the MERFISH images using the method described previously^50^. In short, the imputed expression profile of a MERFISH cell was calculated as the weighted average of the expression profiles of its 30 nearest-neighbour anchor cells in the scRNA-seq dataset in the co-embeded PCA space. The weights were based on the distance between the scRNA-seq cells to the MERFISH cell and were calculated by the find_nearest_anchor function from the SeuratIntegration class using default parameters.

We evaluated the validity of the imputation results by comparing them with the gene expression measured by MERFISH and with the previously measured spatial expression patterns in Allen Brain Atlas in situ hybridization data^51^ for the genes included in the MERFISH gene panel, and with the Allen Brain Atlas in situ hybridization data only for the genes not included in the MERFISH gene panel. We performed two correlation analysis for comparing imputation results with the MERFISH measurement results. First, we calculated for each gene the mean expression level in every cluster from the imputation results and the MERFISH-measurement results. We then quantified for each gene the Pearson correlation coefficient between the imputed cluster means and MERFISH-measured cluster means across all clusters. Second, we calculated for each gene the mean expression levels of every imaged FOVs from the imputation results and the MERFISH-measurement results, and then quantified the Pearson correlation coefficient between the imputed FOV means and MERFISH-measured FOV means across all imaged FOVs. The first comparison evaluated how well the relative expression levels of genes in different clusters were recapitulated by the imputation and the second comparison evaluated how well the spatial dependence of gene expression was recapitulated by the imputation.

For the genes not included in the MERFISH, we visually compared the spatial patterns of gene expression determined by imputation with those determined in Allen Brain Atlas in situ hybridization data.

### MERFISH image registration to Common Coordinate Framework (CCF)

Registration of MERFISH data to the Allen Mouse Brain Common Coordinate Framework (CCF) version 3 was performed in a two-step process involving the reconstruction of 2D MERFISH tissue slices to a 3D volumetric image through alignment of DAPI signals in the MERFISH images to the Nissl template images in Allen Reference Atlas, followed by a 3D refinement using landmarks based on cell types with known localizations in the CCF. For the initial reconstruction, we used the DAPI channel of in the MERFISH images of individual brain slices and the Nissl template images in the Allen Reference Atlas, which is aligned to the Allen CCF. For each MERFISH sample from the same animal, brain slices were ordered and rotated to match coronal or sagittal orientation of the CCF. Coronal slices were cropped ∼200 µm past the midline while sagittal slices were cropped at the posterior end of the cerebellum. In each animal, key slices containing recognizable landmarks were used to identify corresponding CCF planes, and all remaining CCF planes were determined by linear interpolation. To aid the registration process, features in the DAPI image were enhanced by highlighting pixels containing cell types that localized to known brain regions (e.g. VLMC cells at the brain surface, ependymal cells in the ventricles, dentate gyrus granular cells, etc.). The corresponding features in the Nissl image were also highlighted using the CCF annotations and/or morphological operations. Finally, each DAPI/Nissl image pair was registered with an affine, and then B-spline transformation using the program Elastix^112^. Each transformation was then applied to the cell positions to find their initial position in CCF space.

In the second alignment step to refine the CCF registration, an additional 3D-3D registration was performed using additional selected cell types from the MERFISH data that are known to be localized to certain brain regions in the CCF. In total, 36 suitable cell types were identified along with their corresponding brain region annotations in the CCF, as well as two level-1space modules (SM_CTX, SM_RSP) that delineated the cells in the isocortex. These selected cell types (or spatial modules) were each randomly assigned an intensity label, and a 3D volumetric image was generated using their initial positions in the CCF space from the first reconstruction step. A second target 3D image was generated but using only the CCF annotations: For each selected cell type, the corresponding brain region annotations in the CCF were assigned the intensity label for that cell type or spatial module, and all other annotated regions were removed. As before, for certain cell types, morphological operations on certain annotations were used to denote the midline, tissue surface or hollow ventricles. Finally, these two 3D images were registered using a B-spline transformation and the cell positions were refined.

After the MERFISH data was registered to the CCF, each MERFISH cell was assigned a 3-dimensional coordinate, (ccfx, ccfy, ccfz), indicating its spatial location in the CCF space, where ccfx indicates the coordinate value along the rostral-caudal direction, ccfy indicates the coordinate value along the dorsal-ventral direction, and ccfz indicates the coordinate value along the lateral-medial direction. Each MERFISH cell was also assigned a brain region annotation ID as defined in the CCF, indicating its brain region identity.

For visualization in individual figures, we presented the MERFISH-imaged cells in the experimental coordinates, but reverse transformed the brain region boundaries defined in the CCF into the experimental coordinates by reversing the above-described MERFISH image-to-CCF transformation.

### Neurotransmitter identities of the neurons

We assigned neurotransmitter identity to the neurons based on their expression of canonical neurotransmitter transporter genes. Specifically, *Slc17a7*, *Slc17a6* and *Slc17a8* were used for glutamatergic neurons, *Slc32a1* for GABAergic neurons, *Slc6a4* for serotonergic neurons, *Slc6a3* for dopaminergic neurons, *Slc18a3* for cholinergic neurons, *Slc6a5* for glycinergic neurons, and *Slc6a2* for noradrenergic neurons. In addition, *Hdc*, which is involved in histamine synthesis, was used to mark the histaminergic neurons. For all these genes, we used an expression threshold of RNA counts per cell n ≥ 2, determined by MERFISH, to assign neurotransmitter identity to individual neurons.

### Spatial-module analysis

We did two rounds of spatial-module analysis to delineate molecularly defined brain regions based on local cell-type composition.

For the first round of spatial module analysis, we defined a local cell-type-composition vector for each cell to characterize its neighbourhood composition of cell types at the subclass level. We began by finding the 50 spatially nearest neighbours for each cell. Because vascular and immune cells are usually randomly distributed across most brain regions, we excluded them from the spatial-module analysis. Then we assigned a weight to each neighbor cell j of a cell i as:

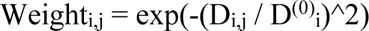

Where D_i,j_ is the spatial distance between cell i and cell j, and D^(0)^_i_ is the distance scaling factor. Because different brain regions have different cell densities, we let D^(0)^_i_ be adjustable based on the local cell density and defined D^(0)^_i_ as two times the distance between cell i and its 5^th^ nearest spatial neighbour. Then we defined the local cell-type-composition vector of a cell from its neighbour cell types and weights. Each element of a local cell-type-composition vector corresponds to a cell type, and the value is the sum of the weights of the spatial neighbours that belong to this cell type.

We generated the first level of spatial modules by clustering cells based on their local cell-type-composition vectors at the subclass level. We normalized the local cell-type-composition vectors by their L2 norms and ran the Leiden clustering method to cluster the cells. We manually curated the clusters by merging the clusters that did not form clear spatial boundaries and annotated the clusters based on the major brain regions that they corresponded to. This round of analysis gave level-1 spatial modules

We then generated the level-2 spatial modules for each level-1 spatial module separately. Because the spatial heterogeneity of cell types within individual major brain regions are mainly due to neurons, we only considered neurons for the second round of spatial module analysis. We calculated the local cell-type-composition vectors using the same method described for the first round of spatial-module analysis with two modifications. The first modification is that we considered both subclasses and clusters to define the local cell-type-composition vectors – the subclass-based vector was concatenated with the cluster-based vector to form the overall vector. The second modification is that we used a shorter distance scaling factor D^(0)^_i_ for the higher spatial resolution in this round. We defined D^(0)^_i_ as the distance between cell i and its 5^th^ nearest spatial neighbour. Then we used the same method described for the first round of spatial module analysis to cluster cells based on their local cell-type-composition vectors to generate level-2 spatial modules.

### Spatial gradient analysis

To define the degree of how discrete or how well separated individual clusters were within each subclass, for each cell we calculated its “neighbourhood purity” defined by the fraction of cells that had the same cell-cluster label as the center cell among its 50 nearest neighbours in the gene-expression space. The discreteness of a cell cluster was defined by the mean value of the neighbourhood purities of all cells within the cluster. We then determined the median cluster discreteness of a subclass as a measure of how discrete individual clusters were within the subclass.

To visualize the spatial gradient of the subclasses or groups of transcriptionally similar subclasses, PCA was used to reduce dimensionality of the normalized expression data, and to calculate a ‘pseudotime’ value for each cell as previously described^8^. Next, spatial gradients were visualized by representing gene expression profiles of the cells using either the first PC (PC1) or the pseudotime value of individual cells on the spatial maps. In addition, correlation of the PC1 or pseudotime values and the spatial coordinate of the cells were plotted. For the IT neurons in the isocortex, cortical depth was used as the spatial coordinate, and was calculated for individual neurons as previously described^8^ for coronal slices in the region between Bregma ∼- 0.8 and ∼+1.7 where the L6b CTX cells formed clear thin layer at the bottom border of isocortex. For the D1 and D2 MSNs, locations along the dorsolateral-ventromedial axis were used as spatial coordinate values and were calculated using the ccfy (dorsal-ventral) and ccfz (medial-lateral) locations of individual cells. For LSX neurons and tanycytes, locations along the dorsal-lateral axis (ccfy) were used as spatial coordinate values.

### Cell-cell interaction analysis

We performed cell-cell interaction analysis at the subclass level. All cells with a subclass label-transfer confidence score greater than 0.8 were used in this analysis. We divided cells into major brain regions based on their CCF coordinates. Due to the high complexity of cell type compositions of hypothalamus, midbrain, and hindbrain, we further divided these regions each into two regions: hypothalamus was divided into anterior and posterior hypothalamus; midbrain was divided into anterior and posterior midbrain; hindbrain was divided into pons and medulla. For hypothalamus and midbrain, the region was divided based on the cell locations along the rostral-caudal axis (ccfx), specifically, the mean value of minimum and maximum ccfx value for all the cells within the region was used to divide the region into anterior and posterior part. We only considered the subclasses that were either enriched or had a sufficient abundance in each brain region for the cell-cell interaction analysis. For neuronal subclasses, we used the enrichment score as described in Figure 2b caption. For anterior hypothalamus, posterior hypothalamus, anterior midbrain, posterior midbrain, pons, and medulla, we used an enrichment score threshold of 6 to stringently select cells in these regions. For the other brain regions, we set the enrichment score threshold to 2. For astrocytes, we used an enrichment score threshold of 1 for all brain regions. For the remaining subclasses of non-neuronal cells, we considered them in a brain region if the total cell number of that subclass was greater than 50 in this region.

For each subclass pair within each region, we determined the number of cell pairs (one from each subclass) that were in contact or proximity and compared the number of contact or proximal cell pairs to a null distribution generated by randomly shifting spatial positions of the cells locally^11^. Two cells were considered in contact or proximity if the distance between the cell centroid positions was within a distance threshold (R_proximal_). We first defined R_proximal_ to be 15 μm, which is comparable to the soma size of the cells in the mouse brain. To generate the null distribution by randomly shifting spatial positions of the cells locally, for each round of randomization, we shifted the spatial location of each cell to a random position within 100 μm from its original location. We did 1000 rounds of randomization. After each round, we calculated the number of cell pairs that were in contact or proximity between every pair of subclasses. For each pair of subclasses, we fitted the distribution of the number of contact/proximal cell pairs generated by 1000 randomizations to a normal distribution to generate the null distribution. We then compared the observed contact/proximal cell-pair number with the null distribution to determine the enrichment fold change and the p-value of the enrichment. Then we used the Benjamini-Hochberg multiple-hypothesis testing correction method to adjust the p-values. We used the adjusted p-value threshold of 0.05 and the number of observed proximal pair threshold of 20 to select pair of subclasses that showed significant probability to be in contact or proximity and called these subclass pairs as interacting cell-type pairs.

Since the stringent distance threshold, R_proximal_ = 15 μm, may eliminate some cell-type pairs that communicate through paracrine signaling, we also relaxed this distance threshold to a greater value (R_proximal_ = 30 um), but for cell-type pairs identified with this relaxed distance threshold, we further required that at least one ligand-receptor pair was upregulated in the proximal cell pairs as compared to non-proximal cell pairs (see below) in order to call these cell types as interacting cell-type pairs.

### Ligand-receptor analysis and analysis of other genes upregulating in interacting cell pairs

We performed the ligand-receptor analysis at the subclass level. All cells with subclass label-transfer confidence score greater than 0.8 were used in this analysis. We used the CellChat database^113^ to define the ligand-receptor pairs. For a ligand-receptor pair k, we defined the ligand-receptor expression score for a pair of cells i and j as:

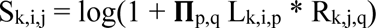

Where L_k,i,p_ is the expression level of the p^th^ component of the ligand of the ligand-receptor pair k in the cell i; R_k,j,q_ is the expression level of the q^th^ component of the receptor of the ligand-receptor pair k in the cell j. The expression levels used here were the imputed gene expression results as described in the “Imputation of transcriptome-wide gene expressions of individual cells in MERFISH images” section.

We performed ligand-receptor pair analysis for the cell-type pairs that showed statistically significant proximity compared to the null distribution as described in the “Cell-cell interaction analysis” section above, using the R_proximal_ = 30 μm. For a pair of cell types and a ligand-receptor pair, we calculated the distributions of ligand-receptor expression scores for all proximal cell pairs, i.e., cell pairs with soma centroid distance smaller than R_proximal_, from this cell-type pair (one cell from each cell type). Then we randomly selected the same number of cell pairs from this cell-type pair with soma centroid distance greater than R_proximal_. We calculated the distributions of ligand-receptor expression scores for the non-proximal cell pairs. We used one-sided Welch’s t-test to test if the mean ligand-receptor expression scores were significantly higher in proximal cell pairs than the scores in the non-proximal cell pairs. Then we used the Benjamini-Hochberg multiple-hypothesis testing correction method to adjust the p-values. We selected significant ligand-receptor pairs that satisfied the following three criteria: the mean of ligand-receptor expression score was at least 2-fold higher in the proximal cell pairs than those in the non-proximal cell pairs; the adjusted p-value was < 0.01; the ligand-receptor expression scores were greater than zero in at least 40% of the proximal cell pairs. Using this approach, we determined the ligand-receptor pairs that were statistically significantly upregulated in the proximal cell pairs as compared to the non-proximal cell pairs in each cell-type pair that showed statistically significant proximity using R_proximal_ = 30.

We then used a similar approach to determine other genes that were upregulated in the proximal cell pairs as compared to non-proximal cell pairs in each cell-type pair. We first determined the highly variable genes for each cell type. Only highly variable genes were considered for this gene upregulation analysis. For each cell-type A that showed significant proximity with another cell type B as compared to the null distribution, we divided the type-A cells into two groups based on whether they were within R_proximal_ of any type-B cells. We calculated for each gene the expression distributions in the two groups respectively and used one-sided Welch’s t-test to test if the mean expression was significantly higher in the first group than that in the second group. We used the Benjamini-Hochberg multiple-hypothesis testing correction method to adjust the p-values. We selected significantly upregulated genes using the following criteria: the mean expression level was at least 2-fold higher in the proximal cell pairs than those in the non-proximal cell pairs; the adjusted p-values were < 0.01.

